## Supplementary material for "Evolution of C4 photosynthesis predicted by constraint-based modelling": environment.yml

```
name: base
channels:
  - gurobi
  - bioconda
  - conda-forge
  - defaults
  - http://conda.anaconda.org/gurobi
dependencies:
  - plotly=3.8.0
  - seaborn=0.7.1
  - pytest=2.9.2
  - python=2.7.*
  - pip==19.1
  - tqdm=4.11.2
  - xlrd=1.0.0
  - goatools=0.6.10
  - numpy==1.13.*
  - scipy==0.19.*
  - pandas==0.20.*
  - pip:
    - cobra==0.8.2
    - colorlover==0.2.1
    - escher==1.6.0
    - optlang==1.2.1
    - tabulate==0.7.7
prefix: //anaconda
```
