## Supplementary material for "Evolution of C4 photosynthesis predicted by constraint-based modelling": 2019-05-06-mb-genC3-CO2-Effect.html


### Effect of the CO2 uptake rate on C3 metabolism¶

#### 0. Initialization¶

##### 0.1 Load Packages¶

In [1]:

```
#Import sys
import sys 
sys.path.append("../src/") 

#Import init for initialisation & loading user-defined functions
from init_fba import *

#Initialize notebook settings
theNotebook = '2019-05-06-mb-genC3-CO2-Effect'
init_notebook(theNotebook)

#load sbml model
c3_model = load_sbml_model()

#inf  
inf = float(1e6)
```

```
//anaconda/lib/python2.7/site-packages/cryptography/hazmat/primitives/constant_time.py:26 CryptographyDeprecationWarning: Support for your Python version is deprecated. The next version of cryptography will remove support. Please upgrade to a 2.7.x release that supports hmac.compare_digest as soon as possible.
```

#### 1. C3 Model¶

##### 1.1 Constraints¶

In [2]:

```
#CONSTRAINT: Set flux of all export reaction to zero
for r_obj in c3_model.reactions:
    r_id = r_obj.id
    if r_id[0:2] == "Ex":
        r_obj.bounds = (0.,0.)

#CONSTRAINT: Divergent fluxes of export and import reactions
set_bounds('Im_CO2', (-inf, inf), c3_model)
set_bounds('Im_H2O', (-inf, inf), c3_model)
set_bounds('Im_H2S', (0.,0.), c3_model)
set_bounds('Im_NH4', (0., 0.), c3_model)
set_bounds('Im_NO3', (0., inf), c3_model)
set_bounds('Im_Pi', (0., inf), c3_model)
set_bounds('Im_SO4', (0., inf), c3_model)
set_bounds('Ex_O2', (-inf, inf), c3_model)
set_bounds('Ex_Suc', (0., inf), c3_model)
set_bounds('Ex_starch', (0., inf), c3_model)
set_bounds('Ex_AA', (0., inf), c3_model)

#CONSTRAINT: 
set_bounds('G6PDH_h', (0.,0.), c3_model)
set_bounds('PPIF6PK_c', (0,0.), c3_model)

#CONSTRAINT: max. photon consumption 1000 μE
set_bounds('Im_hnu', (0, 1000), c3_model)

#CONSTRAINT: CO2 uptake rate in C3 plants is about 20 μmol/(m2*s)
f_CO2 = 20 #[μmol/(m2*s)] 
set_bounds('Im_CO2', (0, f_CO2), c3_model)
```

In [3]:

```
#CONSTRAINT: Maintenace cost

atp_cost_L3_m = 0.009111187245501572 #Mitochondria-L3-ATP Cost [µmol*s-1*m-2]
atp_cost_L3_h = 0.15270708327974447 #Chloroplast-L3-ATP Cost [µmol*s-1*m-2]
atp_cost_L3_p = 0.0076669066992201855 #Peroxisome-L3-ATP Cost [µmol*s-1*m-2]
atp_cost_L3_c = 0.042683072918274702 #Cytosl/Other-L3-ATP Cost [µmol*s-1*m-2]

set_fixed_flux('NGAM_c',atp_cost_L3_c + atp_cost_L3_p, c3_model)
set_fixed_flux('NGAM_m',atp_cost_L3_m, c3_model)
set_fixed_flux('NGAM_h',atp_cost_L3_h, c3_model)
```

In [4]:

```
#CONSTRAINT: Output of sucrose : total amino acid and sucrose : starch
set_fixed_flux_ratio({'Ex_Suc':2.2,'Ex_AA':1.0}, c3_model)
set_fixed_flux_ratio({'Ex_Suc':1.0,'Ex_starch':1.0}, c3_model)
```

Out[4]:

```
<optlang.glpk_interface.Constraint at 0x11892edd0>
```

In [5]:

```
#CONSTRAINT: oxygenation : decarboxylation = 1 : 10
set_fixed_flux_ratio({'RBC_h':10,'RBO_h':1}, c3_model)
```

Out[5]:

```
<optlang.glpk_interface.Constraint at 0x10672c990>
```

In [6]:

```
#CONSTRAINT: fluxes through the chloroplastic NADPH dehydrogenase and plastoquinol oxidase were set to zero 
#because the contributions of NADPH dehydrogenase (Yamamoto et al., 2011) and plastoquinol oxidase 
#(Josse et al., 2000) to photosynthesis are thought to be minor.
set_bounds('AOX4_h',(0,0), c3_model)
set_bounds('iCitDHNADP_h',(0,0), c3_model)
```

In [7]:

```
#CONSTRAINT: NTT is only active at night
set_fixed_flux('Tr_NTT',0, c3_model)
```

In [8]:

```
#CONSTRAINT: No uncoupled pyruvate transport
set_bounds('Tr_Pyr1',(0,0), c3_model)
set_bounds('Tr_Pyr2',(0,0), c3_model)
```

#### 2. FBA¶

In [9]:

```
#Set FBA solver
c3_model.solver = "glpk"

#Optimize/Maximize sucrose output
result_fba_c3 = c3_model.optimize('maximize') #perform FBA
Ex_Suc = c3_model.reactions.get_by_id("Ex_Suc")
Ex_Suc.objective_coefficient = 1.

#Optimize/Minimize total flux
if result_fba_c3.status == 'optimal': 
    result_pfba_c3 = cobra.flux_analysis.parsimonious.pfba(c3_model)

#Fetch flux for CO2 uptake
flux_CO2 = result_pfba_c3.fluxes['Im_CO2']


#Array defining proprtion of CO2 uptake 
L_CO2 = np.linspace(0,1,21)

#Initialize dictionary to store results
D_fba = {}

#Iterate over proportions of CO2 uptake
for v_co2 in L_CO2:
    
    #Fix upper flux bound for photon uptake
    set_bounds('Im_CO2', (0, flux_CO2 * v_co2), c3_model)
    
    #Optimize/Maximize sucrose output
    result_fba_c3 = c3_model.optimize('maximize') #perform FBA
    
    #Optimize/Minimize total flux
    if result_fba_c3.status == 'optimal': # check if feasible
        result_pfba_c3  = cobra.flux_analysis.parsimonious.pfba(c3_model) #perform pFBA
        D_fba[flux_CO2 * v_co2] = result_pfba_c3.fluxes
    else:
        D_fba[flux_CO2 * v_co2] = pd.Series(pd.Series( index=[r_obj.id for r_obj in c3_model.reactions], data = [0]*len(c3_model.reactions)))
```

#### 4 Figures¶

In [10]:

```
xaxis_title = 'CO2 Uptake [µmol/s/m2]'
save_fig = False
```

In [11]:

```
L_r = ['Ex_Suc','Ex_AA']
create_scatter_plot_rxn_c3(D_fba, L_r, 'Phloem Export', xaxis_title, save_fig = save_fig)
```

In [12]:

```
L_r = ['PSI_h','PSII_h']
create_bar_plot_met_c3(c3_model, D_fba, L_r, 'hnu_h', 'Photon Uptake by Photosystems', xaxis_title, save_fig = save_fig)
```

In [13]:

```
L_r = ['NDH1_h','NDH2_h']
create_scatter_plot_rxn_c3(D_fba, L_r, 'Cyclic Electron Flow', xaxis_title, save_fig = save_fig)
```
