## Supplementary material for "Evolution of C4 photosynthesis predicted by constraint-based modelling": 2019-05-06-mb-genC3-Light-Effect.html


### Effect of the PPFD on C3 metabolism¶

#### 0. Initialization¶

In [1]:

```
#Import sys
import sys 
sys.path.append("../src/") 

#Import init for initialisation & loading user-defined functions
from init_fba import *

#Initialize notebook settings
theNotebook = '2019-05-06-mb-genC3-Light-Effect'
init_notebook(theNotebook)

#load sbml model
c3_model = load_sbml_model()

#inf  
inf = float(1e6)
```

In [7]:

```
#CONSTRAINT: NTT is only active at night
set_fixed_flux('Tr_NTT',0, c3_model)
```

In [8]:

```
#CONSTRAINT: No uncoupled pyruvate transport
set_bounds('Tr_Pyr1',(0,0), c3_model)
set_bounds('Tr_Pyr2',(0,0), c3_model)
```

#### 2. FBA¶

In [9]:

```
#Set FBA solver
c3_model.solver = "glpk"

#Optimize/Maximize sucrose output
result_fba_c3 = c3_model.optimize('maximize') #perform FBA
Ex_Suc = c3_model.reactions.get_by_id("Ex_Suc")
Ex_Suc.objective_coefficient = 1.

#Optimize/Minimize total flux
if result_fba_c3.status == 'optimal': 
    result_pfba_c3 = cobra.flux_analysis.parsimonious.pfba(c3_model)

#Fetch flux for photon uptake
flux_hnu = result_pfba_c3.fluxes['Im_hnu']

#Array defining proprtion of light uptake 
L_hnu = np.linspace(0,2,21)

#Initialize dictionary to store results
D_fba = {}

#Iterate over proportions of photon uptake
for hnu in L_hnu:
    
    #Fix flux for photon uptake
    set_bounds('Im_hnu', (flux_hnu * hnu, flux_hnu * hnu), c3_model)
    
    #Optimize/Maximize sucrose output
    result_fba_c3 = c3_model.optimize('maximize') #perform FBA
    
    #Optimize/Minimize total flux
    if result_fba_c3.status == 'optimal': # check if feasible
        result_pfba_c3  = cobra.flux_analysis.parsimonious.pfba(c3_model) #perform pFBA
        D_fba[hnu*flux_hnu] = result_pfba_c3.fluxes
    else:
        D_fba[hnu*flux_hnu] = pd.Series(pd.Series( index=[r_obj.id for r_obj in c3_model.reactions], data = [0]*len(c3_model.reactions)))
```

```
//anaconda/lib/python2.7/site-packages/cobra/util/solver.py:419: UserWarning:

solver status is 'infeasible'
```

#### 4 Figures¶

In [10]:

```
xaxis_title = 'PPFD [µE]'
save_fig = False
```

In [11]:

```
L_r = ['Ex_Suc','Ex_AA']
create_scatter_plot_rxn_c3(D_fba, L_r, 'Phloem Export', xaxis_title, save_fig = save_fig)
```

In [12]:

```
L_r = ['Im_CO2']
create_scatter_plot_rxn_c3(D_fba, L_r, 'CO2 Uptake', xaxis_title, save_fig = save_fig)
```

In [13]:

```
L_r = ['PSI_h','PSII_h']
create_bar_plot_met_c3(c3_model, D_fba, L_r, 'hnu_h', 'Photon Uptake by Photosystems', xaxis_title, save_fig = save_fig)
```
