## Supplementary material for "Evolution of C4 photosynthesis predicted by constraint-based modelling": 2019-05-06-mb-genC3.html


### Input, output fluxes and flux of energy in C3 metabolism¶

#### 0. Initialization¶

In [1]:

```
#Import sys
import sys 
sys.path.append("../src/") 

#Import init for initialisation & loading user-defined functions
from init_fba import *

#Initialize notebook settings
theNotebook = '2019-05-06-mb-genC3'
init_notebook(theNotebook)

#load sbml model
c3_model = load_sbml_model()

#goatools
from goatools import obo_parser
goDB = obo_parser.GODag('../src/go_basic.obo')
```

```
//anaconda/lib/python2.7/site-packages/cryptography/hazmat/primitives/constant_time.py:26 CryptographyDeprecationWarning: Support for your Python version is deprecated. The next version of cryptography will remove support. Please upgrade to a 2.7.x release that supports hmac.compare_digest as soon as possible.
```

```
load obo file ../src/go_basic.obo
../src/go_basic.obo: fmt(1.2) rel(2017-10-20) 49,155 GO Terms
```

##### 0.1 Functions¶

In [2]:

```
def proportion_E_consumption(L_m, name, save_fig = True):
    D_E_Con_RXN = {}
    L_r_E_Con = []


    for m_id in L_m:
        for r_obj in c3_model.metabolites.get_by_id(m_id).reactions:
            r_id = r_obj.id
            if not r_id[:2]  in ['Tr','Ex','Im'] :
                flux = round(result_pfba_c3.fluxes[r_id] * r_obj.get_coefficient(m_id),5)
                if flux < 0 :
                    L_r_E_Con.append(r_id)
                    D_E_Con_RXN[r_id] = abs(flux)

    E_Con_Total = sum(D_E_Con_RXN.values())

    D_E_Con_goTerm = {}
    for r_id in D_E_Con_RXN:
        r_obj = c3_model.reactions.get_by_id(r_id)
        L_goId = r_obj.annotation['go']
        if not isinstance(L_goId, list):
            L_goId = [L_goId]
        L_goTerm = [goDB[goId].name for goId in L_goId]
        if not L_goTerm[0] in D_E_Con_goTerm:
            D_E_Con_goTerm[L_goTerm[0]] = 0
        D_E_Con_goTerm[L_goTerm[0]] += D_E_Con_RXN[r_id]


    D_E_Con_goTerm = {goTerm: flux for goTerm, flux in D_E_Con_goTerm.items() if flux/E_Con_Total > 0.01}
    D_E_Con_goTerm['Others'] = E_Con_Total - sum(D_E_Con_goTerm.values())

    trace = go.Pie(
        labels = D_E_Con_goTerm.keys(),
        values = D_E_Con_goTerm.values(),
        textfont=dict(size=18,family='Arial',),
        marker=dict(colors=[D_goTerm_col[goTerm] for goTerm in D_E_Con_goTerm],
                    line=dict(color='#FFF', width=1)),
    )

    data = [trace]

    layout = go.Layout(
                height=600, 
                width=750,
                title='Proportion of %s Consumption' %name
        )

    fig = go.Figure(data=data, layout=layout)
    
    if save_fig:
        iplot(fig,filename='%s_consumption' %name, image='svg',image_height=500,image_width=750)
        sleep(5)
    else:
        iplot(fig)
    return E_Con_Total
```

In [3]:

```
def proportion_E_production(L_m, name, savefig = True):


    D_E_Pro_RXN = {}
    L_r_E_Pro = []


    for m_id in L_m:
        for r_obj in c3_model.metabolites.get_by_id(m_id).reactions:
            r_id = r_obj.id
            if not r_id[:2]  in ['Tr','Ex','Im'] :
                flux = round(result_pfba_c3.fluxes[r_id] * r_obj.get_coefficient(m_id),5)
                if flux > 0 :
                    L_r_E_Pro.append(r_id)
                    D_E_Pro_RXN[r_id] = abs(flux)

    E_Pro_Total = sum(D_E_Pro_RXN.values())

    D_E_Pro_goTerm = {}
    for r_id in D_E_Pro_RXN:
        r_obj = c3_model.reactions.get_by_id(r_id)
        L_goId = r_obj.annotation['go']
        if not isinstance(L_goId, list):
            L_goId = [L_goId]
        L_goTerm = [goDB[goId].name for goId in L_goId]
        if not L_goTerm[0] in D_E_Pro_goTerm:
            D_E_Pro_goTerm[L_goTerm[0]] = 0
        D_E_Pro_goTerm[L_goTerm[0]] += D_E_Pro_RXN[r_id]


    D_E_Pro_goTerm = {goTerm: flux for goTerm, flux in D_E_Pro_goTerm.items() if flux/E_Pro_Total > 0.01}
    D_E_Pro_goTerm['Others'] = E_Pro_Total - sum(D_E_Pro_goTerm.values())

    trace = go.Pie(
        labels = D_E_Pro_goTerm.keys(),
        values = D_E_Pro_goTerm.values(),
        textfont=dict(size=18,family='Arial',),
        marker=dict(colors=[D_goTerm_col[goTerm] for goTerm in D_E_Pro_goTerm],
                    line=dict(color='#FFF', width=1)),
    )


    data = [trace]

    layout = go.Layout(
                height=600, 
                width=750,
                title='Proportion of %s Production' %name
        )

    fig = go.Figure(data=data, layout=layout)
    
    if save_fig:
        iplot(fig,filename='%s_Production' %name,image='svg',image_height=500,image_width=750)
        sleep(5)
    else:
        iplot(fig)
    return E_Pro_Total
```

#### 1. C3 Model¶

##### 1.1 Constraints¶

In [4]:

```
#CONSTRAINT: Set flux of all export reaction to zero
for r_obj in c3_model.reactions:
    r_id = r_obj.id
    if r_id[0:2] == "Ex":
        r_obj.bounds = (0.,0.)

#CONSTRAINT: Divergent fluxes of export and import reactions
set_bounds('Im_CO2', (-inf, inf), c3_model)
set_bounds('Im_H2O', (-inf, inf), c3_model)
set_bounds('Im_H2S', (0.,0.), c3_model)
set_bounds('Im_NH4', (0., 0.), c3_model)
set_bounds('Im_NO3', (0., inf), c3_model)
set_bounds('Im_Pi', (0., inf), c3_model)
set_bounds('Im_SO4', (0., inf), c3_model)
set_bounds('Ex_O2', (-inf, inf), c3_model)
set_bounds('Ex_Suc', (0., inf), c3_model)
set_bounds('Ex_starch', (0., inf), c3_model)
set_bounds('Ex_AA', (0., inf), c3_model)

#CONSTRAINT: ???
set_bounds('G6PDH_h', (0.,0.), c3_model)
set_bounds('PPIF6PK_c', (0,0.), c3_model)

#CONSTRAINT: max. photon consumption 1000 μE
set_bounds('Im_hnu', (0, 1000), c3_model)

#CONSTRAINT: CO2 uptake rate in C3 plants is about 20 μmol/(m2*s)
f_CO2 = 20 #[μmol/(m2*s)] 
set_bounds('Im_CO2', (0, f_CO2), c3_model)
```

In [5]:

```
#CONSTRAINT: Maintenace cost

atp_cost_L3_m = 0.009111187245501572 #Mitochondria-L3-ATP Cost [µmol*s-1*m-2]
atp_cost_L3_h = 0.15270708327974447 #Chloroplast-L3-ATP Cost [µmol*s-1*m-2]
atp_cost_L3_p = 0.0076669066992201855 #Peroxisome-L3-ATP Cost [µmol*s-1*m-2]
atp_cost_L3_c = 0.042683072918274702 #Cytosl/Other-L3-ATP Cost [µmol*s-1*m-2]

set_fixed_flux('NGAM_c',atp_cost_L3_c + atp_cost_L3_p, c3_model)
set_fixed_flux('NGAM_m',atp_cost_L3_m, c3_model)
set_fixed_flux('NGAM_h',atp_cost_L3_h, c3_model)
```

In [6]:

```
#CONSTRAINT: Output of sucrose : total amino acid and sucrose : starch
set_fixed_flux_ratio({'Ex_Suc':2.2,'Ex_AA':1.0}, c3_model)
set_fixed_flux_ratio({'Ex_Suc':1.0,'Ex_starch':1.0}, c3_model)
```

Out[6]:

```
<optlang.glpk_interface.Constraint at 0x1067d18d0>
```

In [7]:

```
#CONSTRAINT: oxygenation : decarboxylation = 1 : 10
set_fixed_flux_ratio({'RBC_h':10,'RBO_h':1}, c3_model)
```

Out[7]:

```
<optlang.glpk_interface.Constraint at 0x1067d88d0>
```

In [8]:

```
#CONSTRAINT: fluxes through the chloroplastic NADPH dehydrogenase and plastoquinol oxidase were set to zero 
#because the contributions of NADPH dehydrogenase (Yamamoto et al., 2011) and plastoquinol oxidase 
#(Josse et al., 2000) to photosynthesis are thought to be minor.
set_bounds('AOX4_h',(0,0), c3_model)
set_bounds('iCitDHNADP_h',(0,0), c3_model)
```

In [9]:

```
#CONSTRAINT: NTT is only active at night
set_fixed_flux('Tr_NTT',0, c3_model)
```

In [10]:

```
#CONSTRAINT: No uncoupled pyruvate transport
set_bounds('Tr_Pyr1',(0,0), c3_model)
set_bounds('Tr_Pyr2',(0,0), c3_model)
```

#### 2. FBA¶

In [11]:

```
#Set FBA solver
c3_model.solver = "glpk"

#Optimize/Maximize sucrose output
Ex_Suc = c3_model.reactions.get_by_id("Ex_Suc")
Ex_Suc.objective_coefficient = 1.
result_fba_c3 = c3_model.optimize('maximize') #perform FBA


#Optimize/Minimize total flux
if result_fba_c3.status == 'optimal': # check if feasible
    result_pfba_c3 = cobra.flux_analysis.parsimonious.pfba(c3_model) #perform pFBA
```

#### 4 Figures¶

In [12]:

```
save_fig = False
```

In [13]:

```
#Map GO terms to reactions consuming or producing ATP, NADH, NADPH in all compartments

L_m = ['ATP_c', 'ATP_h', 'ATP_m', 'NADH_c', 'NADH_p', 'NADH_m', 'NADH_h', 'NADPH_c', 'NADPH_m', 'NADPH_h']

L_goTerm = []

for m_id in L_m:
    for r_obj in c3_model.metabolites.get_by_id(m_id).reactions:
        r_id = r_obj.id
        if not r_id[:2]  in ['Tr','Ex','Im'] :
            L_goId_r = r_obj.annotation['go']
            if not isinstance(L_goId_r, list):
                L_goId_r = [L_goId_r]
            L_goTerm_r = [goDB[goId].name for goId in L_goId_r]
            L_goTerm += L_goTerm_r

L_goTerm = list(set(L_goTerm))
L_col = ['hsl('+str(h)+',50%'+',50%)' for h in np.linspace(0, 360, len(L_goTerm)+1)]
D_goTerm_col = {L_goTerm[n]: col for n, col in enumerate(L_col) if n < len(L_goTerm)}
D_goTerm_col['Others'] = L_col[-1]
```

In [14]:

```
#Proportion of subsystems in ATP production and consumption
L_m = ['ATP_c', 'ATP_h', 'ATP_m']
ATP_Pro = proportion_E_production(L_m, 'ATP', save_fig)
ATP_Con = proportion_E_consumption(L_m, 'ATP', save_fig)
```

In [15]:

```
#Proportion of subsystems in NADPH production and consumption
L_m = ['NADPH_c', 'NADPH_h', 'NADPH_m']
NADPH_Pro = proportion_E_production(L_m, 'NADPH', save_fig)
NADPH_Con = proportion_E_consumption(L_m, 'NADPH', save_fig)
```

In [16]:

```
#Proportion of subsystems in NADH production and consumption
L_m = ['NADH_c', 'NADH_h', 'NADH_m']
NADH_Pro = proportion_E_production(L_m, 'NADH', save_fig)
NADH_Con = proportion_E_consumption(L_m, 'NADH', save_fig)
```

In [17]:

```
#Proportion of engergy eqivalents

trace = go.Pie(
    labels = ['ATP', 'NADPH', 'NADH'],
    values = [ATP_Pro, NADPH_Pro, NADH_Pro],
    textfont=dict(size=18,family='Arial'),
    marker=dict(line=dict(color='#FFF', width=1)),
)

data = [trace]

layout = go.Layout(
            height=500, 
            width=750, 
            title = 'Proportion of Energy Equivalents'
    )

fig = go.Figure(data=data, layout=layout)

if save_fig:
    iplot(fig,filename='E_Pro_proportion',image='svg',image_height=500,image_width=750)
    sleep(5)
else:
    iplot(fig)
```

In [18]:

```
#Proportion of maintenace costs on respiratory ATP

L_r_NGAM = ['NGAM_h','NGAM_c','NGAM_m']
L_flux = [result_pfba_c3.fluxes[r_id] for r_id in L_r_NGAM]
print(L_flux)
cplx5_m = c3_model.reactions.get_by_id('cplx5_m')
ATP_cplx5 = result_pfba_c3.fluxes['cplx5_m'] * cplx5_m.get_coefficient('ATP_m')

trace = go.Pie(
    labels = L_r_NGAM + ['Others'],
    values = L_flux + [ATP_cplx5 - sum(L_flux)],
    textfont=dict(size=18,family='Arial',),
    marker=dict(line=dict(color='#FFF', width=1)),
)

data = [trace]

layout = go.Layout(
            title="Proportion of Respiratory ATP",
            height=500, 
            width=750,
            
    )

fig = go.Figure(data=data, layout=layout)

if save_fig:
    iplot(fig,filename='maintenance_respiratory_ATP',image='svg',image_height=500,image_width=750)
    sleep(5)
else:
    iplot(fig)
    
print('ATP Maintainence : Respiratory ATP  =  %s%%' %(round(sum(L_flux) / (ATP_cplx5) * 100,3)))
```

```
[0.15270708327974447, 0.050349979617494885, 0.009111187245501572]
```

```
ATP Maintainence : Respiratory ATP  =  28.092%
```

In [19]:

```
#save results to excel
df = pd.DataFrame(columns=['rxn','subsystem','flux'])
L_subsystem = []
L_transport = []
for r_obj in c3_model.reactions:
    L_goId = r_obj.annotation['go']
    if not isinstance(L_goId, list):
        L_goId = [L_goId]
    L_goTerm = [goDB[goId].name for goId in L_goId]
    L_subsystem = L_subsystem + L_goTerm
    if 'GO:0006810' in L_goId:
        L_transport.append(r_obj.id)
    df.loc[r_obj.id] = [r_obj.reaction,', '.join(L_goTerm),result_pfba_c3.fluxes[r_obj.id]]

writer = pd.ExcelWriter(theNotebook+'/excel/flux_solution.xlsx')
df.to_excel(writer)
writer.save()
```

In [20]:

```
#Print characteristic numbers of model
print('Number of reactions: %s' %len(c3_model.reactions))
L_r_transport = [r_id for r_id in L_transport if r_id[:2] == 'Tr']
L_r_export = [r_id for r_id in L_transport if r_id[:2] == 'Ex']
L_r_import = [r_id for r_id in L_transport if r_id[:2] == 'Im']
print('Number of internal transporters: %s' %len(L_r_transport))
print('Number of export reactions: %s' %len(L_r_export))
print('Number of import reactions: %s' %len(L_r_import))
print('Number of subsystems: %s' %len(list(set(L_subsystem))))
print('Number of metabolites: %s' %len(c3_model.metabolites))
```

```
Number of reactions: 572
Number of internal transporters: 139
Number of export reactions: 90
Number of import reactions: 8
Number of subsystems: 59
Number of metabolites: 413
```

In [21]:

```
#Create dataframe for input and output reactions
df = pd.DataFrame(columns=['RXN','Flux'])
for r_id in L_r_export:
    r_flux = result_pfba_c3.fluxes[r_id]
    if round(r_flux,5) != 0:
        r_obj = c3_model.reactions.get_by_id(r_id)
        df.loc[r_id] = [r_obj.reaction,r_flux]
for r_id in L_r_import:
    r_flux = result_pfba_c3.fluxes[r_id]
    if abs(round(r_flux,5)) > 0:
        r_obj = c3_model.reactions.get_by_id(r_id)
        df.loc[r_id] = [r_obj.reaction,r_flux]

max_flux = max(df['Flux'])
min_flux = min(df['Flux'])
```

In [22]:

```
#Create figure for input and output reactions
i_max = 3
j_max = 3
r_max = len(df.index)
trace = {}
D_r_name = {}

for n, r_id in enumerate(df.index):
    D_r_name[n+1] = r_id
    r_flux = df.get_value(r_id, 'Flux')
    trace[n+1] = go.Bar(
        x=[1],
        y=[r_flux],
        text = [str(round(r_flux,3))],
        textposition= 'auto',
        showlegend=False,
        marker=dict(
                color='rgba(219, 64, 82, 0.7)' if r_id[:2] == 'Ex' else 'rgba(50, 171, 96, 0.7)',
                line=dict(
                        color='rgba(219, 64, 82, 1.0)' if r_id[:2] == 'Ex' else 'rgba(50, 171, 96, 1.0)',
                        width=2)))

fig = ply.tools.make_subplots(rows=i_max, cols=j_max, subplot_titles=D_r_name.values())

k = 1
for i in range(1,i_max+1):
    for j in range(1,j_max+1):
        if k <= r_max:
            fig.append_trace(trace[k], i, j)
        else:
            break
        k += 1
        
fig['layout'].update(height=1000, width=1000)


for k in range(1,i_max*j_max+1):
    if k <= r_max:
            fig['layout']['yaxis'+str(k)].update(title='Flux  [µmol/s/m2]', range=[0, max_flux])
            fig['layout']['xaxis'+str(k)].update(showticklabels=False)
    else:
            break

if save_fig:
    iplot(fig, filename='input_output_fluxes',image='svg',image_height=1000,image_width=1000)
    sleep(5)
else:
    iplot(fig)
```

```
This is the format of your plot grid:
[ (1,1) x1,y1 ]  [ (1,2) x2,y2 ]  [ (1,3) x3,y3 ]
[ (2,1) x4,y4 ]  [ (2,2) x5,y5 ]  [ (2,3) x6,y6 ]
[ (3,1) x7,y7 ]  [ (3,2) x8,y8 ]  [ (3,3) x9,y9 ]
```
