## Supplementary material for "Evolution of C4 photosynthesis predicted by constraint-based modelling": 2019-05-06-mb-genC4-C4-mode.html


### C4 GenC4 Model - Effect of C4 mode on the emergence of the C4 cycle¶

#### 0. Initialization¶

In [1]:

```
#Import sys
import sys 
sys.path.append("../src/") 

#Import init for initialisation & loading user-defined functions
from init_fba import *

#Initialize notebook settings
theNotebook = '2019-05-06-mb-genC4-Decarb-Enzyme-Effect'
init_notebook(theNotebook)

#load sbml model
c3_model = load_sbml_model()
```

```
//anaconda/lib/python2.7/site-packages/cryptography/hazmat/primitives/constant_time.py:26 CryptographyDeprecationWarning: Support for your Python version is deprecated. The next version of cryptography will remove support. Please upgrade to a 2.7.x release that supports hmac.compare_digest as soon as possible.
```

#### 1. C3 Model¶

##### 1.1 Constraints¶

In [2]:

```
#CONSTRAINT: Set flux of all export reaction to zero
for r_obj in c3_model.reactions:
    r_id = r_obj.id
    if r_id[0:2] == "Ex":
        r_obj.bounds = (0.,0.)

#CONSTRAINT: Divergent fluxes of export and import reactions
set_bounds('Im_CO2', (-inf, inf), c3_model)
set_bounds('Im_H2O', (-inf, inf), c3_model)
set_bounds('Im_H2S', (0.,0.), c3_model)
set_bounds('Im_NH4', (0., 0.), c3_model)
set_bounds('Im_NO3', (0., inf), c3_model)
set_bounds('Im_Pi', (0., inf), c3_model)
set_bounds('Im_SO4', (0., inf), c3_model)
set_bounds('Ex_O2', (-inf, inf), c3_model)
set_bounds('Ex_Suc', (0., inf), c3_model)
set_bounds('Ex_starch', (0., inf), c3_model)
set_bounds('Ex_AA', (0., inf), c3_model)

#CONSTRAINT: 
set_bounds('G6PDH_h', (0.,0.), c3_model)
set_bounds('PPIF6PK_c', (0,0.), c3_model)

#CONSTRAINT: max. photon consumption auf C4 plants 1000 μE
set_bounds('Im_hnu', (0, 1000), c3_model)

#CONSTRAINT: CO2 uptake rate uin C4 plants is higher, about 40 μmol/(m2*s)
f_CO2 = 40 #[μmol/(m2*s)] 
set_bounds('Im_CO2', (0, f_CO2), c3_model)
```

In [3]:

```
#CONSTRAINT: Maintenace cost

atp_cost_L3_m = 0.009111187245501572 #Mitochondria-L3-ATP Cost [µmol*s-1*m-2]
atp_cost_L3_h = 0.15270708327974447 #Chloroplast-L3-ATP Cost [µmol*s-1*m-2]
atp_cost_L3_p = 0.0076669066992201855 #Peroxisome-L3-ATP Cost [µmol*s-1*m-2]
atp_cost_L3_c = 0.042683072918274702 #Cytosl/Other-L3-ATP Cost [µmol*s-1*m-2]

set_fixed_flux('NGAM_c',atp_cost_L3_c + atp_cost_L3_p, c3_model)
set_fixed_flux('NGAM_m',atp_cost_L3_m, c3_model)
set_fixed_flux('NGAM_h',atp_cost_L3_h, c3_model)
```

In [4]:

```
#CONSTRAINT: fluxes through the chloroplastic NADPH dehydrogenase and plastoquinol oxidase were set to zero 
#because the contributions of NADPH dehydrogenase (Yamamoto et al., 2011) and plastoquinol oxidase 
#(Josse et al., 2000) to photosynthesis are thought to be minor.
set_bounds('AOX4_h',(0,0), c3_model)
set_bounds('iCitDHNADP_h',(0,0), c3_model)
```

In [5]:

```
#CONSTRAINT: NTT is only active at night
set_fixed_flux('Tr_NTT',0, c3_model)
```

In [6]:

```
#CONSTRAINT: No uncoupled pyruvate transport
set_bounds('Tr_Pyr1',(0,0), c3_model)
set_bounds('Tr_Pyr2',(0,0), c3_model)
```

#### 2. C4 Model¶

##### 2.1 Compose Model¶

###### 2.1.1 Two copies of genC3 model and exchange reactions¶

In [7]:

```
c4_model = cobra.Model('c4_model')

cell_types = ['M', 'B']

#duplicate metabolites
for m in c3_model.metabolites:
    for cell in cell_types:
        m_dt = cobra.Metabolite('['+cell+']_'+m.id, name = m.formula, compartment = m.compartment)
        c4_model.add_metabolites([m_dt])

#duplicate reactions
for r_c3_obj in c3_model.reactions:
    for cell in cell_types:
        r_c4_obj = cobra.Reaction('['+cell+']_'+r_c3_obj.id)
        r_c4_obj.name = r_c3_obj.name
        r_c4_obj.subsystem = r_c3_obj.subsystem
        r_c4_obj.bounds = r_c3_obj.bounds
        c4_model.add_reaction(r_c4_obj)
        r_c4_obj.add_metabolites({'['+cell+']_'+m_c3_obj.id: r_c3_obj.get_coefficient(m_c3_obj) for m_c3_obj in r_c3_obj.metabolites})
        
        
#metabolites excluded from M/BS exchange
no_transport = ['NO3','NO2', 'O2','Na', 'H2S', 'SO4',
                'H2O','FBP','F26BP','DPGA','H','ACD','AC','M_DASH_THF', '5M_DASH_THF', 'H_DASH_Cys', 'aH_DASH_Cys', 'ORO', 'DHO',
                'GABA','A_DASH_Ser','PRPP','AD','THF','DHF','ADN','Mas','CoA','GluP',
                'A_DASH_CoA','cellulose1','cellulose2','cellulose3','starch1',
                'starch2','starch3','TRXox','TRXrd','Glu_DASH_SeA','T6P','aMet',
                'PPi', 'P5C', 'NH4', 'Pi', 'CO2', 'OAA','HCO3', 
                'UTP', 'UDP', 'UDPG', 'ATP', 'ADP', 'AMP', 'IMP', 'XMP', 
                'GTP', 'GDP', 'GMP', 'OMP', 'UMP', 'CTP', 'GDP', 'CDP', 'dADP', 
                'dCDP', 'dGDP', 'dUDP', 'dUTP', 'dUMP', 'dTMP', 'dTDP', 'GTP', 
                'dATP', 'dCTP', 'dGTP', 'dTTP', 'NAD', 'NADH', 'NADP', 'NADPH']

#add M/BS exchange reactions
L_r_transport = []
for m_c3_obj in c3_model.metabolites:
    if m_c3_obj.id[-1:] == 'c' and m_c3_obj.id[:-2] not in no_transport:
        r_c4_obj = cobra.Reaction('[MB]_'+m_c3_obj.id)
        r_c4_obj.name = '[MB]_'+m_c3_obj.id
        r_c4_obj.subsystem = 'Exchange'
        r_c4_obj.bounds = (-inf, inf)
        c4_model.add_reaction(r_c4_obj)
        r_c4_obj.add_metabolites({'[M]_'+m_c3_obj.id: -1,'[B]_'+m_c3_obj.id: 1 })
        L_r_transport.append('[MB]_'+m_c3_obj.id)
```

###### 2.1.2 Adaptations for second unconstrained rubisco population in the bundle sheath¶

In [8]:

```
#CONSTRAINT: Add external CO2 species to bundle sheath
#(the original CO2 species is treated as internal CO2)
m_list_CO_Ex= ['[B]_CO2_ex_c','[B]_CO2_ex_h']

for m_id in m_list_CO_Ex:
    m_obj = cobra.Metabolite(m_id)
    c4_model.add_metabolites(m_obj)

#CONSTRAINT: Copy reactions 'Tr_CO2h', 'RBC_h' and replace internal CO2 with external CO2 in the copied reactions 
r_list_CO_Ex = ['Tr_CO2h', 'RBC_h']

for r_id in r_list_CO_Ex:
    r_obj = c4_model.reactions.get_by_id('[B]_'+r_id)
    r_obj_Ex = cobra.Reaction(r_obj.id+'_Ex')
    r_obj_Ex.name = r_obj.id+'_Ex'
    r_obj_Ex.subsystem = r_obj.subsystem
    r_obj_Ex.bounds = r_obj.bounds
    c4_model.add_reaction(r_obj_Ex)
    r_obj_Ex.add_metabolites({m_obj.id if not m_obj.id[:-2] == '[B]_CO2' else '[B]_CO2_ex'+m_obj.id[-2:]: r_obj.get_coefficient(m_obj) 
                                  for m_obj in r_obj.metabolites})

#CONSTRAINT: CO2 exchange between mesophyll and bundle sheat
r_c4_obj = cobra.Reaction('[MB]_CO2_c')
r_c4_obj.name = '[MB]_CO2_c'
r_c4_obj.subsystem = 'Exchange'
r_c4_obj.bounds = (-inf, inf)
c4_model.add_reaction(r_c4_obj)
r_c4_obj.add_metabolites({'[M]_CO2_c': -1,'[B]_CO2_ex_c': 1 })
L_r_transport.append('[MB]_CO2_c')
```

##### 2.3 Constraints¶

In [9]:

```
#CONSTRAINT: No CO2 uptake in bundle sheat cells due to suberin layer in cell membranes
set_fixed_flux('[B]_Im_CO2',0, c4_model)
```

In [10]:

```
#CONSTRAINT: Output of sucrose : total amino acid and sucrose : starch
set_fixed_flux_ratio({'[B]_Ex_Suc':2.2,'[B]_Ex_AA':1.0}, c4_model)
set_fixed_flux_ratio({'[B]_Ex_Suc':1.0,'[B]_Ex_starch':1.0}, c4_model)

set_fixed_flux_ratio({'[M]_Ex_Suc':2.2,'[M]_Ex_AA':1.0}, c4_model)
set_fixed_flux_ratio({'[M]_Ex_Suc':1.0,'[M]_Ex_starch':1.0}, c4_model)
```

Out[10]:

```
<optlang.glpk_interface.Constraint at 0x117b46550>
```

In [11]:

```
#Reaction variables for light uptake
B_Im_hnu = c4_model.reactions.get_by_id("[B]_Im_hnu")
M_Im_hnu = c4_model.reactions.get_by_id("[M]_Im_hnu")

#CONSTRAINT: Total Photon uptake limited to 1000 µE
const_hnu_sum = c4_model.problem.Constraint( B_Im_hnu.flux_expression + M_Im_hnu.flux_expression,
                                        lb = 0, ub = 1000)

c4_model.add_cons_vars(const_hnu_sum)

#CONSTRAINT: Total Photon uptake by bundle sheath must be less equal than in mesophyll
const_hnu_ratio = c4_model.problem.Constraint( M_Im_hnu.flux_expression - B_Im_hnu.flux_expression,
                                        lb = 0, ub = 1000)

c4_model.add_cons_vars(const_hnu_ratio)
```

In [12]:

```
#CONSTRAINT: oxygenation : carboxylation = 1 : 3
set_fixed_flux_ratio({'[B]_RBC_h_Ex':1,'[B]_RBO_h':1},c4_model)
set_fixed_flux_ratio({'[M]_RBC_h':1,'[M]_RBO_h':1},c4_model)
```

Out[12]:

```
<optlang.glpk_interface.Constraint at 0x118155390>
```

#### 3 FBA¶

In [13]:

```
#Define experiments, here: the three decraboxylation enzymes PEP-CK (in model: [B]_PEPC1_c), NAD-ME (in model: [B]_MalDH2_m) 
#and NADP-ME (in model: [B]_MalDH4_h)
D_exp = {'[B]_PEPC1_c': 'PEP-CK', '[B]_MalDH2_m': 'NAD-ME', '[B]_MalDH4_h': 'NADP-ME'}

#Set FBA solver
c4_model.solver = "glpk"

#Initialize dictionaries to store results
D_fba = {}
D_pfba = {}
D_fva = {}
D_pfva = {}

#Reaction Variables
B_Ex_Suc = c4_model.reactions.get_by_id("[B]_Ex_Suc")
B_RBO = c4_model.reactions.get_by_id("[B]_RBO_h")
M_RBO = c4_model.reactions.get_by_id("[M]_RBO_h")

#Set flux reactions related to decraboxylation enzymes to zero 
for exp in D_exp:
    set_fixed_flux(exp,0, c4_model)
            
#Run every FBA experiment
for exp in D_exp:
    
    #Allow non-zero flux for current decarboxylation enzyme
    set_bounds(exp, (0,inf), c4_model)           
    
    #Optimize/Maximize sucrose output
    B_Ex_Suc.objective_coefficient = 1.
    c4_model_copy = c4_model.copy()
    result_fba = c4_model_copy.optimize('maximize')
    del c4_model_copy
    c4_model.objective =[]
    set_fixed_flux(B_Ex_Suc.id,result_fba.fluxes[B_Ex_Suc.id], c4_model)
    
    #Optimize/Minimize oxygenation rate by rubisco (set True)
    if True:
            B_RBO.objective_coefficient = 1.
            M_RBO.objective_coefficient = 1.
            c4_model_copy = c4_model.copy()
            result_fba = c4_model_copy.optimize('minimize')
            del c4_model_copy
            c4_model.objective =[]
            set_fixed_flux(B_RBO.id,result_fba.fluxes[B_RBO.id],c4_model)            
            set_fixed_flux(M_RBO.id,result_fba.fluxes[M_RBO.id],c4_model)    

    D_fba[exp] = result_fba.fluxes            #store FBA results
    
    #Optimize/Minimize total flux and perform (p)FVA (set True)
    if True: 
        if result_fba.status == 'optimal':
            c4_model_copy = c4_model.copy()
            #perform pFBA
            result_pfba = cobra.flux_analysis.parsimonious.pfba(c4_model_copy)
            D_pfba[exp] = result_pfba.fluxes     
            D_pfba[exp]= D_pfba[exp].set_value('total',result_pfba.f)
            #perform FVA
            result_fva = cobra.flux_analysis.flux_variability_analysis(c4_model_copy)     #perform FVA
            D_fva[exp] = result_fva  
            #perform pFVA results with 1.5% deviation of pFBA total flux
            result_pfva = cobra.flux_analysis.flux_variability_analysis(c4_model_copy,pfba_factor=1.015) 
            D_pfva[exp] = result_pfva 
            del c4_model_copy      
    
    #Reset flux of current decarboxylation enzyme to zero
    set_fixed_flux(exp,0, c4_model)           
    
    #Reset reaction bounds
    set_bounds(B_RBO.id,(0,inf),c4_model)
    set_bounds(M_RBO.id,(0,inf),c4_model)
    set_bounds(B_Ex_Suc.id,(0,inf),c4_model)
```

In [14]:

```
save_fba_to_excel(c3_model, c4_model, D_pfba, D_exp, theNotebook)
save_fva_to_excel(c3_model, c4_model, D_pfva, D_exp, theNotebook)
```

In [15]:

```
for exp, exp_name in D_exp.items():
    save_to_json(exp_name, D_pfba[exp], theNotebook)
```

#### 4 Figures¶

In [16]:

```
xaxis_title = 'decarboxylation enzyme'
save_fig = False
```

In [17]:

```
create_bar_plot_rxn(D_pfba, D_exp,
                    {'total':'Total',},
                    'Total Flux', xaxis_title,
                    stacked = True, save_fig=save_fig)
```

In [18]:

```
create_bar_plot_rxn(D_pfba, D_exp,
                    {'[B]_Ex_Suc':'Ex_Suc',},
                    'Sucrose Export', xaxis_title,
                    stacked = True, save_fig=save_fig)
```

In [19]:

```
create_bar_plot_rxn(D_pfba, D_exp,
                    {'[M]_Im_hnu':'Mesophyll','[B]_Im_hnu': 'Bundle sheath'},
                    'Photon Uptake', xaxis_title,
                    stacked = True, save_fig=save_fig)
```

In [20]:

```
create_bar_plot_met(D_pfba, D_exp,
                    {'[M]_PSII_h':'Mesophyll','[B]_PSII_h': 'Bundle sheath'},'hnu_h',
                    'Photon Uptake by PSII', xaxis_title, c3_model,
                    stacked = True, save_fig=save_fig)
```

In [21]:

```
create_bar_plot_met(D_pfba, D_exp,
                    {'[M]_PSI_h':'Mesophyll','[B]_PSI_h': 'Bundle sheath'},'hnu_h',
                    'Photon Uptake by PSI', xaxis_title, c3_model,
                    stacked = True, save_fig=save_fig)
```

In [22]:

```
create_bar_plot_met(D_pfba, D_exp,
                    {'[M]_cplx5_m':'Mesophyll','[B]_cplx5_m': 'Bundle sheath'},'ATP_m',
                    'ATP Synthesis Mitochondria', xaxis_title, c3_model,
                    stacked = True, save_fig=save_fig)
```

In [23]:

```
create_bar_plot_met(D_pfba, D_exp,
                    {'[M]_ATPase_h':'Mesophyll','[B]_ATPase_h': 'Bundle sheath'},'ATP_h',
                    'ATP Synthesis Chloroplast', xaxis_title, c3_model,
                    stacked = True, save_fig=save_fig)
```

In [24]:

```
create_bar_plot_rxn(D_pfba, D_exp,
                    {'[M]_RBC_h':'Mesophyll (constrained)','[B]_RBC_h_Ex':'Bundle sheat (constrained)',
                 '[B]_RBC_h': 'Bundle sheat (unconstrained)'},
                    'Carboxylation by Rubisco',  xaxis_title,
                    stacked = True, save_fig=save_fig)
```

In [25]:

```
create_bar_plot_rxn(D_pfba, D_exp,
                    {'[M]_PEPC2_c': 'Mesophyll', '[B]_PEPC2_c': 'Bundle sheath'},
                    'PEPC',  xaxis_title,
                     stacked = True, save_fig=save_fig)
```

In [26]:

```
create_bar_plot_rxn(D_pfba, D_exp,
                    {'[M]_PyrPiDK_h': 'Mesophyll', '[B]_PyrPiDK_h': 'Bundle sheath'},
                    'PyrPiDK',  xaxis_title,
                     stacked = True, save_fig=save_fig)
```

In [27]:

```
create_bar_plot_rxn(D_fba, D_exp,
                    {'[M]_iCitDHNADP_h': 'Mesophyll', '[B]_iCitDHNADP_h': 'Bundle sheath'},
                    'iCitDHNADP_h', 'oxygenation : decarboxylation ratio',
                     stacked = True, save_fig=save_fig)
```

In [28]:

```
create_bar_plot_rxn(D_fba, D_exp,
                    {'[M]_AOX4_h': 'Mesophyll', '[B]_AOX4_h': 'Bundle sheath'},
                    'AOX4_h', 'oxygenation : decarboxylation ratio',
                     stacked = True, save_fig=save_fig)
```

In [29]:

```
D_comp = {'c':'cytsol','h':'chloroplast','m':'mitochondrium'}

D_comp_rxn = {}
L_max_flux = []
for comp in D_comp:
    m_obj = c4_model.metabolites.get_by_id('[B]_CO2_'+comp)
    L_rxn = []
    for r_obj in m_obj.reactions:
        if not r_obj.id[4:6] in ['Tr','Im']:
            lb = r_obj.bounds[0]
            v = r_obj.get_coefficient(m_obj.id)
            if (lb >= 0 and v>0) or (lb < 0 and v<0):  
                L_rxn.append(r_obj.id)
                L_max_flux = L_max_flux + [v*D_pfba[x][r_obj.id] for x in D_exp]
    D_comp_rxn[comp] = L_rxn
max_flux =  max(L_max_flux)

for comp, L_rxn in D_comp_rxn.items():
    create_bar_plot_rxn(D_pfba, D_exp,
                        L_rxn,
                        'CO2 Production Bundesheat (%s)' %(D_comp[comp]),  xaxis_title,
                        stacked = True, save_fig=save_fig)
```

In [30]:

```
plot_transport(D_pfba, D_exp, L_r_transport, xaxis_title, save_fig=save_fig)
```

In [31]:

```
L_r_index = ['[MB]_Mal_c','[MB]_Pyr_c','[MB]_Asp_c','[MB]_Ala_c', 
             '[MB]_PGA_c', '[MB]_2PGA_c', '[MB]_GAP_c', '[MB]_DHAP_c',
             '[MB]_Suc_c', '[MB]_Frc_c', '[MB]_F6P_c', '[MB]_G6P_c', '[MB]_G1P_c', '[MB]_S6P_c',
             '[MB]_Glu_c',
             '[MB]_PEP_c', '[MB]_KG_c'
            ]

plot_transport_fva(c4_model, D_exp, D_pfva, D_pfba, L_r_transport, 40, 'FVA of Exchange Reaction (pFBA Factor = 1.5%)', L_r_index=L_r_index, save_fig=save_fig)
```

In [32]:

```
#Display snippet of the flux distribution using Escher map
for exp, exp_name in D_exp.items():
    display(HTML('<h1>%s</h1>' %exp_name))
    b = Builder(map_json='../data/2018-06-29-mb-C4-Map-Decarb-Enzymes.json',reaction_data=D_pfba[exp].to_dict())
    display(b.display_in_notebook(height=1000))
```

### NADP-ME

### NAD-ME

### PEP-CK
