## Supplementary material for "Evolution of C4 photosynthesis predicted by constraint-based modelling": 2019-05-06-mb-genC4-CO2-Limitation.html


### Effect of CO2 limitation on the C4 mode¶

#### 0. Initialization¶

In [1]:

```
#Import sys
import sys 
sys.path.append("../src/") 

#Import init for initialisation & loading user-defined functions
from init_fba import *

#Initialize notebook settings
theNotebook = '2019-05-06-mb-genC4-CO2-Limitation'
init_notebook(theNotebook)

#load sbml model
c3_model = load_sbml_model()
```

###### 2.1.2 Adaptations for second unconstrained rubisco population in the bundle sheath¶

In [8]:

```
#CONSTRAINT: Add external CO2 species in bundle sheat
#(the original CO2 species is treated as internal CO2)
m_list_CO_Ex= ['[B]_CO2_ex_c','[B]_CO2_ex_h','[B]_CO2_ex_m']

for m_id in m_list_CO_Ex:
    m_obj = cobra.Metabolite(m_id)
    c4_model.add_metabolites(m_obj)

#CONSTRAINT: Copy all reactions using internal CO2 and exchange internal with external CO2 in the copied recactions
r_list_CO_Ex = ['Tr_CO2h', 'RBC_h']

for r_id in r_list_CO_Ex:
    r_obj = c4_model.reactions.get_by_id('[B]_'+r_id)
    r_obj_Ex = cobra.Reaction(r_obj.id+'_Ex')
    r_obj_Ex.name = r_obj.id+'_Ex'
    r_obj_Ex.subsystem = r_obj.subsystem
    r_obj_Ex.bounds = r_obj.bounds
    c4_model.add_reaction(r_obj_Ex)
    r_obj_Ex.add_metabolites({m_obj.id if not m_obj.id[:-2] == '[B]_CO2' else '[B]_CO2_ex'+m_obj.id[-2:]: r_obj.get_coefficient(m_obj) 
                                  for m_obj in r_obj.metabolites})

#CONSTRAINT: CO2 exchange between mesophyll and bundle sheat
r_c4_obj = cobra.Reaction('[MB]_CO2_c')
r_c4_obj.name = '[MB]_CO2_c'
r_c4_obj.subsystem = 'Exchange'
r_c4_obj.bounds = (-inf, inf)
c4_model.add_reaction(r_c4_obj)
r_c4_obj.add_metabolites({'[M]_CO2_c': -1,'[B]_CO2_ex_c': 1 })
L_r_transport.append('[MB]_CO2_c')
```

##### 2.3 Constraints¶

In [9]:

```
#CONSTRAINT: No CO2 uptake in bundle sheat cells due to suberin layer in cell membranes
set_fixed_flux('[B]_Im_CO2',0, c4_model)
```

In [10]:

```
#CONSTRAINT: Output of sucrose : total amino acid and sucrose : starch
set_fixed_flux_ratio({'[B]_Ex_Suc':2.2,'[B]_Ex_AA':1.0}, c4_model)
set_fixed_flux_ratio({'[B]_Ex_Suc':1.0,'[B]_Ex_starch':1.0}, c4_model)

set_fixed_flux_ratio({'[M]_Ex_Suc':2.2,'[M]_Ex_AA':1.0}, c4_model)
set_fixed_flux_ratio({'[M]_Ex_Suc':1.0,'[M]_Ex_starch':1.0}, c4_model)
```

Out[10]:

```
<optlang.glpk_interface.Constraint at 0x106817090>
```

In [11]:

```
#Reaction variables for light uptake
B_Im_hnu = c4_model.reactions.get_by_id("[B]_Im_hnu")
M_Im_hnu = c4_model.reactions.get_by_id("[M]_Im_hnu")

#CONSTRAINT: Total Photon uptake limited to 1000 µE
const_hnu_sum = c4_model.problem.Constraint( B_Im_hnu.flux_expression + M_Im_hnu.flux_expression,
                                        lb = 0, ub = 1000)

c4_model.add_cons_vars(const_hnu_sum)

#CONSTRAINT: Total Photon uptake by bundle sheath must be less equal than in mesophyll
const_hnu_ratio = c4_model.problem.Constraint( M_Im_hnu.flux_expression - B_Im_hnu.flux_expression,
                                        lb = 0, ub = 1000)

c4_model.add_cons_vars(const_hnu_ratio)
```

In [12]:

```
#CONSTRAINT: oxygenation : carboxylation = 1 : 3
set_fixed_flux_ratio({'[B]_RBC_h_Ex':3,'[B]_RBO_h':1}, c4_model)
set_fixed_flux_ratio({'[M]_RBC_h':3,'[M]_RBO_h':1}, c4_model)
```

Out[12]:

```
<optlang.glpk_interface.Constraint at 0x118441fd0>
```

#### 3 FBA¶

In [13]:

```
#Dictionary defining the experiments according to the CO2-limitation
D_exp = {value: value for value in np.arange(0,41,5)}

#Reaction Variables
B_Ex_Suc = c4_model.reactions.get_by_id("[B]_Ex_Suc")
B_RBO = c4_model.reactions.get_by_id("[B]_RBO_h")
M_RBO = c4_model.reactions.get_by_id("[M]_RBO_h")
M_Im_CO2 = c4_model.reactions.get_by_id("[M]_Im_CO2")

#Set FBA solver
c4_model.solver = "glpk"

#Initialize dictionary to store results
D_fba={}

#Run every FBA experiment

for value in sorted(D_exp.keys()): #iterate over proportions of carboxylation
    
    #Add CO2-limitation constraint 
    const_CO2 = c4_model.problem.Constraint( M_Im_CO2.flux_expression,
                                        lb = value, ub = value)
    c4_model.add_cons_vars(const_CO2)
    
    #Optimize/Maximize sucrose output
    B_Ex_Suc.objective_coefficient = 1.
    c4_model_copy = c4_model.copy()
    result_fba = c4_model_copy.optimize('maximize')
    del c4_model_copy
    c4_model.objective =[]
    set_fixed_flux(B_Ex_Suc.id,result_fba.fluxes[B_Ex_Suc.id], c4_model)
    
    #Optimize/Minimize oxygenation rate by rubisco (set True)
    if True:
            B_RBO.objective_coefficient = 1.
            M_RBO.objective_coefficient = 1.
            c4_model_copy = c4_model.copy()
            result_fba = c4_model_copy.optimize('minimize')
            del c4_model_copy
            c4_model.objective =[]
            set_fixed_flux(B_RBO.id,result_fba.fluxes[B_RBO.id],c4_model)            
            set_fixed_flux(M_RBO.id,result_fba.fluxes[M_RBO.id],c4_model)
    
        
    #Optimize/Minimize total flux
    if result_fba.status == 'optimal': 
        c4_model_copy = c4_model.copy()
        result_pfba = cobra.flux_analysis.parsimonious.pfba(c4_model_copy)
        D_fba[value] = result_pfba.fluxes
        del c4_model_copy 
    
    #Reset reaction bounds
    set_bounds(B_RBO.id,(0,inf),c4_model)
    set_bounds(M_RBO.id,(0,inf),c4_model)
    set_bounds(B_Ex_Suc.id,(0,inf),c4_model)

    #Remove CO2-limitation constraint
    c4_model.remove_cons_vars(const_CO2)
```

#### 4 Figures¶

In [14]:

```
xaxis_title = 'CO2 Uptake [µmol/s/m2]'
save_fig = False
```

In [15]:

```
create_bar_plot_rxn(D_fba, D_exp,
                    {'[M]_Im_hnu':'Mesophyll','[B]_Im_hnu': 'Bundle sheath'},
                    'Photon Uptake', xaxis_title,
                    stacked = True, save_fig=save_fig)
```

In [16]:

```
create_bar_plot_met(D_fba, D_exp,
                    {'[M]_PSII_h':'Mesophyll','[B]_PSII_h': 'Bundle sheath'},'hnu_h',
                    'Photon Uptake by PSII', xaxis_title, c3_model,
                    stacked = True, save_fig=save_fig)
```

In [17]:

```
create_bar_plot_met(D_fba, D_exp,
                    {'[M]_PSI_h':'Mesophyll','[B]_PSI_h': 'Bundle sheath'},'hnu_h',
                    'Photon Uptake by PSI', xaxis_title, c3_model,
                    stacked = True, save_fig=save_fig)
```

In [18]:

```
create_bar_plot_met(D_fba, D_exp,
                    {'[M]_cplx5_m':'Mesophyll','[B]_cplx5_m': 'Bundle sheath'},'ATP_m',
                    'ATP Synthesis Mitochondria', xaxis_title, c3_model,
                    stacked = True, save_fig=save_fig)
```

In [19]:

```
create_bar_plot_met(D_fba, D_exp,
                    {'[M]_ATPase_h':'Mesophyll','[B]_ATPase_h': 'Bundle sheath'},'ATP_h',
                    'ATP Synthesis Chloroplast', xaxis_title, c3_model,
                    stacked = True, save_fig=save_fig)
```

In [20]:

```
create_bar_plot_rxn(D_fba, D_exp,
                    {'[M]_RBC_h':'Mesophyll (constrained)','[B]_RBC_h_Ex':'Bundle sheat (constrained)',
                 '[B]_RBC_h': 'Bundle sheat (unconstrained)'},
                    'Carboxylation by Rubisco', xaxis_title,
                    stacked = True, save_fig=save_fig)
```

In [21]:

```
create_bar_plot_rxn(D_fba, D_exp,
                    {'[B]_MalDH4_h': 'NADP-ME', '[B]_MalDH2_m': 'NAD-ME', '[B]_PEPC1_c': 'PEPCK',},
                    'Decarboxylation Enzymes', xaxis_title,
                    c=True, stacked = True, save_fig=save_fig)
```

In [22]:

```
create_bar_plot_rxn(D_fba, D_exp,
                    {'[M]_PEPC2_c': 'Mesophyll', '[B]_PEPC2_c': 'Bundle sheath'},
                    'PEPC', xaxis_title,
                     stacked = True, save_fig=save_fig)
```

In [23]:

```
create_bar_plot_rxn(D_fba, D_exp,
                    {'[M]_PyrPiDK_h': 'Mesophyll', '[B]_PyrPiDK_h': 'Bundle sheath'},
                    'PyrPiDK', xaxis_title,
                     stacked = True, save_fig=save_fig)
```

In [24]:

```
plot_transport(D_fba, D_exp, L_r_transport, xaxis_title, save_fig=save_fig)
```
