## Supplementary material for "Evolution of C4 photosynthesis predicted by constraint-based modelling": 2019-05-06-mb-genC4-Decarb-Oxy-Ratio-Effect.html


### Analysing the effect of oxygenation : carboxylation ratio on the emergence of the C4 cycle¶

#### 0. Initialization¶

In [1]:

```
#Import sys
import sys 
sys.path.append("../src/") 

#Import init for initialisation & loading user-defined functions
from init_fba import *

#Initialize notebook settings
theNotebook = '2019-05-06-mb-genC4-Decarb-Oxy-Ratio-Effect'
init_notebook(theNotebook)

#load sbml model
c3_model = load_sbml_model()
```

#### 3 FBA¶

In [12]:

```
#Dictionary defining the experiments according to the proportions of carboxylation
D_exp = {ratio: '1 : %s' %ratio for ratio in np.append(np.arange(1,3.25,0.25), np.arange(4,11,1))}

#Reaction Variables
M_Im_hnu = c4_model.reactions.get_by_id("[M]_Im_hnu")
B_Im_hnu = c4_model.reactions.get_by_id("[B]_Im_hnu")
B_Ex_Suc = c4_model.reactions.get_by_id("[B]_Ex_Suc")
B_RBO = c4_model.reactions.get_by_id("[B]_RBO_h")
M_RBO = c4_model.reactions.get_by_id("[M]_RBO_h")

#Set FBA solver
c4_model.solver = "glpk"

#Initialize dictionary to store results
D_fba={}

#Run every FBA experiment

for ratio in sorted(D_exp.keys()): #iterate over proportions of carboxylation
    
    #Add oxygenation:carboxylation constraint 
    const_Rubisco_B = set_fixed_flux_ratio({'[B]_RBC_h_Ex':ratio,'[B]_RBO_h':1},c4_model)
    const_Rubisco_M = set_fixed_flux_ratio({'[M]_RBC_h':ratio,'[M]_RBO_h':1},c4_model)
    
    #Optimize/Maximize sucrose output
    B_Ex_Suc.objective_coefficient = 1.
    c4_model_copy = c4_model.copy()
    result_fba = c4_model_copy.optimize('maximize')
    del c4_model_copy
    c4_model.objective =[]
    set_fixed_flux(B_Ex_Suc.id,result_fba.fluxes[B_Ex_Suc.id], c4_model)
    
    #Optimize/Minimize oxygenation rate by rubisco (set True)
    if False:
            B_RBO.objective_coefficient = 1.
            M_RBO.objective_coefficient = 1.
            c4_model_copy = c4_model.copy()
            result_fba = c4_model_copy.optimize('minimize')
            del c4_model_copy
            c4_model.objective =[]
            set_fixed_flux(B_RBO.id,result_fba.fluxes[B_RBO.id],c4_model)            
            set_fixed_flux(M_RBO.id,result_fba.fluxes[M_RBO.id],c4_model)
    
    #Optimize/Minimize photon uptake (PPFD) (set True)
    if False:
        B_Im_hnu.objective_coefficient = 1.
        M_Im_hnu.objective_coefficient = 1.
        c4_model_copy = c4_model.copy()
        result_fba = c4_model_copy.optimize('minimize')
        del c4_model_copy
        c4_model.objective =[]
        set_fixed_flux(B_Im_hnu.id,result_fba.fluxes[B_Im_hnu.id], c4_model) 
        set_fixed_flux(M_Im_hnu.id,result_fba.fluxes[M_Im_hnu.id], c4_model)
        
    #Optimize/Minimize total flux
    if result_fba.status == 'optimal':      
        c4_model_copy = c4_model.copy()
        result_pfba = cobra.flux_analysis.parsimonious.pfba(c4_model_copy)
        D_fba[ratio] = result_pfba.fluxes
        del c4_model_copy 
    
    #Reset reaction bounds
    set_bounds(B_RBO.id,(0,inf),c4_model)
    set_bounds(M_RBO.id,(0,inf),c4_model)
    set_bounds(B_Ex_Suc.id,(0,inf),c4_model)
    set_bounds(B_Im_hnu.id,(0,1000), c4_model)
    set_bounds(M_Im_hnu.id,(0,1000), c4_model)
    
    #Remove oxygenation:carboxylation constraint
    c4_model.remove_cons_vars(const_Rubisco_B)
    c4_model.remove_cons_vars(const_Rubisco_M)
```

In [13]:

```
save_fba_to_excel(c3_model, c4_model, D_fba, D_exp, theNotebook)
```

In [14]:

```
for exp, exp_name in D_exp.items():
    save_to_json(exp_name, D_fba[exp], theNotebook)
```

#### 4 Figures¶

In [15]:

```
xaxis_title = 'oxygenation : decarboxylation ratio'
save_fig = False
```

In [16]:

```
create_bar_plot_rxn(D_fba, D_exp,
                    {'[M]_Im_hnu':'Mesophyll','[B]_Im_hnu': 'Bundle sheath'},
                    'Photon Uptake', xaxis_title,
                    stacked = True, save_fig=save_fig)
```

In [17]:

```
create_bar_plot_met(D_fba, D_exp,
                    {'[M]_PSII_h':'Mesophyll','[B]_PSII_h': 'Bundle sheath'},'hnu_h',
                    'Photon Uptake by PSII', xaxis_title, c3_model,
                    stacked = True, save_fig=save_fig)
```

In [18]:

```
create_bar_plot_met(D_fba, D_exp,
                    {'[M]_PSI_h':'Mesophyll','[B]_PSI_h': 'Bundle sheath'},'hnu_h',
                    'Photon Uptake by PSI', xaxis_title, c3_model,
                    stacked = True, save_fig=save_fig)
```

In [19]:

```
create_bar_plot_met(D_fba, D_exp,
                    {'[M]_cplx5_m':'Mesophyll','[B]_cplx5_m': 'Bundle sheath'},'ATP_m',
                    'ATP Synthesis Mitochondria', xaxis_title, c3_model,
                    stacked = True, save_fig=save_fig)
```

In [20]:

```
create_bar_plot_met(D_fba, D_exp,
                    {'[M]_ATPase_h':'Mesophyll','[B]_ATPase_h': 'Bundle sheath'},'ATP_h',
                    'ATP Synthesis Chloroplast', xaxis_title, c3_model,
                    stacked = True, save_fig=save_fig)
```

In [21]:

```
create_bar_plot_rxn(D_fba, D_exp,
                    {'[M]_RBC_h':'Mesophyll (constrained)','[B]_RBC_h_Ex':'Bundle sheat (constrained)',
                 '[B]_RBC_h': 'Bundle sheat (unconstrained)'},
                    'Carboxylation by Rubisco', 'oxygenation : decarboxylation ratio',
                    stacked = True, save_fig=save_fig)
```

In [22]:

```
create_bar_plot_rxn(D_fba, D_exp,
                    {'[B]_MalDH4_h': 'NADP-ME', '[B]_MalDH2_m': 'NAD-ME', '[B]_PEPC1_c': 'PEPCK', '[B]_GlyDH_m': 'Gly DH'},
                    'Decarboxylation Enzymes', 'oxygenation : decarboxylation ratio',
                    c=True, stacked = True, save_fig=save_fig)
```

In [23]:

```
4.687/40*100
```

Out[23]:

```
11.7175
```

In [24]:

```
create_bar_plot_rxn(D_fba, D_exp,
                    {'[M]_PEPC2_c': 'Mesophyll', '[B]_PEPC2_c': 'Bundle sheath'},
                    'PEPC', 'oxygenation : decarboxylation ratio',
                     stacked = True, save_fig=save_fig)
```

In [25]:

```
create_bar_plot_rxn(D_fba, D_exp,
                    {'[M]_PyrPiDK_h': 'Mesophyll', '[B]_PyrPiDK_h': 'Bundle sheath'},
                    'PyrPiDK', 'oxygenation : decarboxylation ratio',
                     stacked = True, save_fig=save_fig)
```

In [26]:

```
plot_transport(D_fba, D_exp, L_r_transport, xaxis_title, save_fig=save_fig)
```

In [27]:

```
#Display snippet of the flux distribution using Escher map
for exp, exp_name in D_exp.items():
    if exp in [1,3,10]:
        display(HTML('<h1>%s</h1>' %exp_name))
        b = Builder(map_json='../data/2018-06-29-mb-C4-RBO-RBC-Ratio.json',reaction_data=D_fba[exp].to_dict())
        b.set_reaction_styles(['color','size'])
        display(b.display_in_notebook(height=1000))
```

# 1 : 1.0

# 1 : 3.0

# 1 : 10.0
