## Supplementary material for "Evolution of C4 photosynthesis predicted by constraint-based modelling": 2019-05-06-mb-genC4-H2O-Limitation.html


### Effect of H2O limitation on the C4 mode¶

#### 0. Initialization¶

In [1]:

```
#Import sys
import sys 
sys.path.append("../src/") 

#Import init for initialisation & loading user-defined functions
from init_fba import *

#Initialize notebook settings
theNotebook = '2018-05-06-mb-genC4-H2O-Limitation'
init_notebook(theNotebook)

#load sbml model
c3_model = load_sbml_model()
```

In [12]:

```
#CONSTRAINT: oxygenation : carboxylation = 1 : 3
set_fixed_flux_ratio({'[B]_RBC_h_Ex':3,'[B]_RBO_h':1}, c4_model)
set_fixed_flux_ratio({'[M]_RBC_h':3,'[M]_RBO_h':1}, c4_model)
```

Out[12]:

```
<optlang.glpk_interface.Constraint at 0x106650b50>
```

#### 3 FBA¶

In [13]:

```
#Dictionary defining the experiments according to the H2O-limitation
D_exp = {value: value for value in np.arange(0,41,5)}

#Reaction Variables
B_Ex_Suc = c4_model.reactions.get_by_id("[B]_Ex_Suc")
B_RBO = c4_model.reactions.get_by_id("[B]_RBO_h")
M_RBO = c4_model.reactions.get_by_id("[M]_RBO_h")
M_Im_H2O = c4_model.reactions.get_by_id("[M]_Im_H2O")
B_Im_H2O = c4_model.reactions.get_by_id("[B]_Im_H2O")

#Set FBA solver
c4_model.solver = "glpk"

#Initialize dictionary to store results
D_fba={}

#Run every FBA experiment

for value in sorted(D_exp.keys()): #iterate over proportions of carboxylation
    
    #Add H2O-limitation constraint 
    const_H2O = c4_model.problem.Constraint( M_Im_H2O.flux_expression + B_Im_H2O.flux_expression,
                                        lb = 0, ub = value)
    c4_model.add_cons_vars(const_H2O)
    
    #Optimize/Maximize sucrose output
    B_Ex_Suc.objective_coefficient = 1.
    c4_model_copy = c4_model.copy()
    result_fba = c4_model_copy.optimize('maximize')
    del c4_model_copy
    c4_model.objective =[]
    set_fixed_flux(B_Ex_Suc.id,result_fba.fluxes[B_Ex_Suc.id], c4_model)
    
    #Optimize/Minimize oxygenation rate by rubisco (set True)
    if True:
            B_RBO.objective_coefficient = 1.
            M_RBO.objective_coefficient = 1.
            c4_model_copy = c4_model.copy()
            result_fba = c4_model_copy.optimize('minimize')
            del c4_model_copy
            c4_model.objective =[]
            set_fixed_flux(B_RBO.id,result_fba.fluxes[B_RBO.id],c4_model)            
            set_fixed_flux(M_RBO.id,result_fba.fluxes[M_RBO.id],c4_model)
    
        
    #Optimize/Minimize total flux
    if result_fba.status == 'optimal': 
        c4_model_copy = c4_model.copy()
        result_pfba = cobra.flux_analysis.parsimonious.pfba(c4_model_copy)
        D_fba[value] = result_pfba.fluxes
        del c4_model_copy 
    
    #Reset reaction bounds
    set_bounds(B_RBO.id,(0,inf),c4_model)
    set_bounds(M_RBO.id,(0,inf),c4_model)
    set_bounds(B_Ex_Suc.id,(0,inf),c4_model)

    #Remove H2O-limitation constraint 
    c4_model.remove_cons_vars(const_H2O)
```

#### 4 Figures¶

In [14]:

```
xaxis_title = 'H2O Uptake [µmol/s/m2]'
save_fig = False
```

In [15]:

```
create_bar_plot_rxn(D_fba, D_exp,
                    {'[M]_Im_hnu':'Mesophyll','[B]_Im_hnu': 'Bundle sheath'},
                    'Photon Uptake', xaxis_title,
                    stacked = True, save_fig=save_fig)
```

In [16]:
