## Supplementary material for "Evolution of C4 photosynthesis predicted by constraint-based modelling": 2019-05-06-mb-genC4-Light-Effect.html


### Effect of light on the C4 mode¶

#### 0. Initialization¶

In [1]:

```
#Import sys
import sys 
sys.path.append("../src/") 

#Import init for initialisation & loading user-defined functions
from init_fba import *

#Initialize notebook settings
theNotebook = '2019-05-06-mb-genC4-Light-Effect'
init_notebook(theNotebook)

#load sbml model
c3_model = load_sbml_model()
```

###### 2.1.1 Two copies of genC3 model and exchange reactions¶

In [7]:

```
c4_model = cobra.Model('c4_model')

cell_types = ['M', 'B']

#duplicate metabolites
for m in c3_model.metabolites:
    for cell in cell_types:
        m_dt = cobra.Metabolite('['+cell+']_'+m.id, name = m.formula, compartment = m.compartment)
        c4_model.add_metabolites([m_dt])

#duplicate reactions
for r_c3_obj in c3_model.reactions:
    for cell in cell_types:
        r_c4_obj = cobra.Reaction('['+cell+']_'+r_c3_obj.id)
        r_c4_obj.name = r_c3_obj.name
        r_c4_obj.subsystem = r_c3_obj.subsystem
        r_c4_obj.bounds = r_c3_obj.bounds
        c4_model.add_reaction(r_c4_obj)
        r_c4_obj.add_metabolites({'['+cell+']_'+m_c3_obj.id: r_c3_obj.get_coefficient(m_c3_obj) for m_c3_obj in r_c3_obj.metabolites})
        
        
#metabolites excluded from M/BS exchange
no_transport = ['NO3','NO2', 'O2','Na', 'H2S', 'SO4',
                'H2O','FBP','F26BP','DPGA','H','ACD','AC','M_DASH_THF', '5M_DASH_THF', 'H_DASH_Cys', 'aH_DASH_Cys', 'ORO', 'DHO',
                'GABA','A_DASH_Ser','PRPP','AD','THF','DHF','ADN','Mas','CoA','GluP',
                'A_DASH_CoA','cellulose1','cellulose2','cellulose3','starch1',
                'starch2','starch3','TRXox','TRXrd','Glu_DASH_SeA','T6P','aMet',
                'PPi', 'P5C', 'NH4', 'Pi', 'CO2', 'OAA','HCO3', 
                'UTP', 'UDP', 'UDPG', 'ATP', 'ADP', 'AMP', 'IMP', 'XMP', 
                'GTP', 'GDP', 'GMP', 'OMP', 'UMP', 'CTP', 'GDP', 'CDP', 'dADP', 
                'dCDP', 'dGDP', 'dUDP', 'dUTP', 'dUMP', 'dTMP', 'dTDP', 'GTP', 
                'dATP', 'dCTP', 'dGTP', 'dTTP', 'NAD', 'NADH', 'NADP', 'NADPH']

#add M/BS exchange reactions
R_metTrans = []
for m_c3_obj in c3_model.metabolites:
    if m_c3_obj.id[-1:] == 'c' and m_c3_obj.id[:-2] not in no_transport:
        r_c4_obj = cobra.Reaction('[MB]_'+m_c3_obj.id)
        r_c4_obj.name = '[MB]_'+m_c3_obj.id
        r_c4_obj.subsystem = 'Exchange'
        r_c4_obj.bounds = (-inf, inf)
        c4_model.add_reaction(r_c4_obj)
        r_c4_obj.add_metabolites({'[M]_'+m_c3_obj.id: -1,'[B]_'+m_c3_obj.id: 1 })
        R_metTrans.append('[MB]_'+m_c3_obj.id)
```

###### 2.1.2 Adaptations for second unconstrained rubisco population in the bundle sheath¶

In [8]:

```
#CONSTRAINT: Add external CO2 species in bundle sheat
#(the original CO2 species is treated as internal CO2)
m_list_CO_Ex= ['[B]_CO2_ex_c','[B]_CO2_ex_h','[B]_CO2_ex_m']

for m_id in m_list_CO_Ex:
    m_obj = cobra.Metabolite(m_id)
    c4_model.add_metabolites(m_obj)

#CONSTRAINT: Copy all reactions using internal CO2 and exchange internal with external CO2 in the copied recactions
r_list_CO_Ex = ['Tr_CO2h', 'RBC_h']


for r_id in r_list_CO_Ex:
    r_obj = c4_model.reactions.get_by_id('[B]_'+r_id)
    r_obj_Ex = cobra.Reaction(r_obj.id+'_Ex')
    r_obj_Ex.name = r_obj.id+'_Ex'
    r_obj_Ex.subsystem = r_obj.subsystem
    r_obj_Ex.bounds = r_obj.bounds
    c4_model.add_reaction(r_obj_Ex)
    r_obj_Ex.add_metabolites({m_obj.id if not m_obj.id[:-2] == '[B]_CO2' else '[B]_CO2_ex'+m_obj.id[-2:]: r_obj.get_coefficient(m_obj) 
                                  for m_obj in r_obj.metabolites})

#CONSTRAINT: CO2 exchange between mesophyll and bundle sheat
r_c4_obj = cobra.Reaction('[MB]_CO2_c')
r_c4_obj.name = '[MB]_CO2_c'
r_c4_obj.subsystem = 'Exchange'
r_c4_obj.bounds = (-inf, inf)
c4_model.add_reaction(r_c4_obj)
r_c4_obj.add_metabolites({'[M]_CO2_c': -1,'[B]_CO2_ex_c': 1 })
R_metTrans.append('[MB]_CO2_c')
```

##### 2.3 Constraints¶

In [9]:

```
#CONSTRAINT: No CO2 uptake in bundle sheat cells due to suberin layer in cell membranes
set_fixed_flux('[B]_Im_CO2',0, c4_model)
```

In [10]:

```
#CONSTRAINT: Output of sucrose : total amino acid and sucrose : starch
set_fixed_flux_ratio({'[B]_Ex_Suc':2.2,'[B]_Ex_AA':1.0}, c4_model)
set_fixed_flux_ratio({'[B]_Ex_Suc':1.0,'[B]_Ex_starch':1.0}, c4_model)

set_fixed_flux_ratio({'[M]_Ex_Suc':2.2,'[M]_Ex_AA':1.0}, c4_model)
set_fixed_flux_ratio({'[M]_Ex_Suc':1.0,'[M]_Ex_starch':1.0}, c4_model)
```

Out[10]:

```
<optlang.glpk_interface.Constraint at 0x11822df90>
```

In [11]:

```
#CONSTRAINT: oxygenation : carboxylation = 1 : 3
set_fixed_flux_ratio({'[B]_RBC_h_Ex':3,'[B]_RBO_h':1}, c4_model)
set_fixed_flux_ratio({'[M]_RBC_h':3,'[M]_RBO_h':1}, c4_model)
```

Out[11]:

```
<optlang.glpk_interface.Constraint at 0x118267f90>
```

#### 3 FBA¶

In [12]:

```
#Dictionary of Decarboxylation enzmes and their model ids
D_DCE = {'[B]_MalDH4_h': 'NADP-ME', '[B]_MalDH2_m': 'NAD-ME', '[B]_PEPC1_c': 'PEP-CK'}

#Reaction Variables
M_Im_hnu = c4_model.reactions.get_by_id("[M]_Im_hnu")
B_Im_hnu = c4_model.reactions.get_by_id("[B]_Im_hnu")
B_Ex_Suc = c4_model.reactions.get_by_id("[B]_Ex_Suc")
B_RBO = c4_model.reactions.get_by_id("[B]_RBO_h")
M_RBO = c4_model.reactions.get_by_id("[M]_RBO_h")

#Array defining porportions of light hitting the mesophyll
L_hnu_M = np.concatenate((np.arange(0.5,1.,0.1),np.arange(1,11,1)),axis = 0)

#Maximum light uptake
hnu_max = 1000

#Array defing total light sums
L_hnu_max = sorted([0,50,150] + range(100,hnu_max+100,100))

#Set FBA solver
c4_model.solver = "glpk"

#Initialize dictionary to store results
D_fba={}

#Run every FBA experiment
#iterate L_hnu_max
for hnu_total in L_hnu_max:                
    
    #Add total light contraint 
    const_hnu1 = c4_model.problem.Constraint( M_Im_hnu.flux_expression + B_Im_hnu.flux_expression, 
                                             lb = hnu_total, ub = hnu_total)
    c4_model.add_cons_vars(const_hnu1)
    
    #initialize dictionary to store results
    D_fba[hnu_total] = {}                
    
    #iterate over L_hnu_M 
    for hnu in L_hnu_M:               
        
        #Add light distribution contraint
        const_hnu2 = set_fixed_flux_ratio({'[M]_Im_hnu':hnu,'[B]_Im_hnu':1},c4_model)
        
        #Optimize/Maximize sucrose output
        B_Ex_Suc.objective_coefficient = 1.
        c4_model_copy = c4_model.copy()
        result_fba = c4_model_copy.optimize('maximize')
        del c4_model_copy
        c4_model.objective =[]
        set_fixed_flux(B_Ex_Suc.id,result_fba.fluxes[B_Ex_Suc.id], c4_model)
        
        #Optimize/Minimize oxygenation rate by rubisco (set True)
        if True:
                B_RBO.objective_coefficient = 1.
                M_RBO.objective_coefficient = 1.
                c4_model_copy = c4_model.copy()
                result_fba = c4_model_copy.optimize('minimize')
                del c4_model_copy
                c4_model.objective =[]
                set_fixed_flux(B_RBO.id,result_fba.fluxes[B_RBO.id],c4_model)            
                set_fixed_flux(M_RBO.id,result_fba.fluxes[M_RBO.id],c4_model)
            
        #Optimize/Minimize total flux
        if result_fba.status == 'optimal':      
            c4_model_copy = c4_model.copy()
            result_pfba = cobra.flux_analysis.parsimonious.pfba(c4_model_copy)
            D_fba[hnu_total][hnu] = result_pfba.fluxes
            del c4_model_copy
        else:
            D_fba[hnu_total][hnu]={r_obj.id: 0 for r_obj in c4_model.reactions}
        
        #Reset reaction bounds
        set_bounds(B_RBO.id,(0,inf),c4_model)
        set_bounds(M_RBO.id,(0,inf),c4_model)
        set_bounds(B_Ex_Suc.id,(0,inf),c4_model)
        
        #Remove light distribution constaint
        c4_model.remove_cons_vars(const_hnu2)  
        
    #Remove total light constraint    
    c4_model.remove_cons_vars(const_hnu1)
```

```
//anaconda/lib/python2.7/site-packages/cobra/util/solver.py:419: UserWarning:

solver status is 'infeasible'
```

#### 4 Figures¶

In [13]:

```
save_fig = False
```

In [15]:

```
L_r = ['[B]_MalDH4_h','[B]_PEPC1_c','[B]_MalDH2_m']

L_CO2 = [np.mean([D_fba[hnu_total][hnu_ratio]['[M]_Im_CO2'] 
              if D_fba[hnu_total][hnu_ratio]['[M]_Im_CO2'] > 0 
              else 0 for hnu_ratio in L_hnu_M]) for hnu_total in L_hnu_max ]

L_CO2_var = [np.var([D_fba[hnu_total][hnu_ratio]['[M]_Im_CO2'] 
              if D_fba[hnu_total][hnu_ratio]['[M]_Im_CO2'] > 0 
              else 0 for hnu_ratio in L_hnu_M]) for hnu_total in L_hnu_max ]
    
L_x_axis = range(1,len(L_hnu_max)+1,1)
L_y_axis = range(1,len(L_hnu_M)+1,1)
    
fig = ply.tools.make_subplots(rows=2, cols=3, shared_yaxes=True, shared_xaxes=False, 
                              specs=[[{'colspan':3}, None, None],[{},{},{}]],
                              subplot_titles=('','CO2 Uptake','','NADP-ME','PEPCK','NAD-ME'))

trace = go.Scatter(
        y= L_CO2,
        x= L_x_axis,
        line = {'color':'rgb(214, 39, 40)'})

fig.append_trace(trace, 1, 1)

for i, r_id in enumerate(L_r):
    M_FBA = [[D_fba[hnu_total][hnu_ratio][r_id]/D_fba[hnu_total][hnu_ratio]['[M]_Im_CO2'] 
              if D_fba[hnu_total][hnu_ratio]['[M]_Im_CO2'] > 0 
              else 0 for hnu_total in L_hnu_max] for hnu_ratio in L_hnu_M]
    
    trace = go.Heatmap(
        z=M_FBA,
        x= L_x_axis,
        y= L_y_axis,
        colorbar = {'title':'Decarboxylation Rate : CO2 Uptake Rate','titleside':'right','y':0.19,'len':0.405},
        showscale = True if i+1 == 3 else False,
        zmin = 0,
        zmax = 1)
    
    
    fig.append_trace(trace, 2, i+1)
    
fig['layout'].update(width=1000, height=800,
    yaxis1 = {'title':'Flux [µmol/s/m2]'},
    xaxis1 = {'title':'Total PPFD [µE]','tickmode':'array','tickvals': L_x_axis, 'ticktext':L_hnu_max, 'tickangle':45},
    xaxis2 = {'scaleanchor':'y','scaleratio':4,'tickmode':'array','tickvals': L_x_axis, 'ticktext':L_hnu_max,  'tickangle':45},
    xaxis3 = {'scaleanchor':'y','scaleratio':4,'title':'Total PPFD [µE]','tickmode':'array','tickvals': L_x_axis, 'ticktext':L_hnu_max, 'tickangle':45},
    xaxis4 = {'scaleanchor':'y','scaleratio':4,'tickmode':'array','tickvals': L_x_axis, 'ticktext':L_hnu_max, 'tickangle':45},
    yaxis2 = {'title':'PPFD (BS) : PPFD (M)', 
                 'tickmode':'array','tickvals': L_y_axis, 'ticktext':['1 : %s' %round(hnu,1) for hnu in L_hnu_M]})

if  save_fig:
    iplot(fig,image='svg', filename=theNotebook+'-Figure-1',image_width=1000,image_height=800)
    sleep(5)
else:
    iplot(fig)
```

```
This is the format of your plot grid:
[ (1,1) x1,y1           -                -      ]
[ (2,1) x2,y2 ]  [ (2,2) x3,y2 ]  [ (2,3) x4,y2 ]
```
