## Supplementary material for "Evolution of C4 photosynthesis predicted by constraint-based modelling": README.md

[![Binder](https://mybinder.org/badge_logo.svg)](https://mybinder.org/v2/gh/ma-blaetke/CBM_C3_C4_Metabolism/master)

# Evolution of C4 photosynthesis predicted by constraint-based modelling 
[Mary-Ann Blätke]<sup>1</sup>, [Andrea Bräutigam]<sup>2</sup>

<sup>1</sup>  Leibniz Institute of Plant Genetics and Crop Plant Research (IPK), Corrensstraße 3, 06466 Stadt Seeland OT Gatersleben, Germany)

<sup>2</sup> Computational Biology, Faculty of Biology, Bielefeld University, Universitätsstraße 25, 33615 Bielefeld, Germany

In this repository, we provide all Figure Source Data for the eLife Manuscript, including addtional code and data to run the jupyter notebooks. 
The used one-cell model and its flux solution can be found in the [data](data) directory.
The directory [notebooks](notebooks) contains all jupyter notebooks. To run the jupyter notebooks on a local machine, please check [environment.yml](environment.yml) for information required packages and versions.Jupyter notebooks can also be run by clicking on the [![Binder](https://mybinder.org/badge_logo.svg)](https://mybinder.org/v2/gh/ma-blaetke/CBM_C3_C4_Metabolism/master) button next to them in the list below. 

### Figure 1
- [Figure 1–source data 1. SBML code of the genC3 model](data/2018-23-05-mb-genC3.sbml)
- [Figure 1–source data 2. Flux solution of one-cell C3 model](data/c3_flux_solution_one_cell_model.csv)
- [Figure 1–source data 3. Jupyter notebook - Predicted Fluxes of C3 metabolism](notebooks/2019-05-06-mb-genC3.ipynb)   [![Binder](https://mybinder.org/badge_logo.svg)](https://mybinder.org/v2/gh/ma-blaetke/CBM_C3_C4_Metabolism/master?filepath=notebooks%2F2019-05-06-mb-genC3.ipynb)
- [Figure 1–source data 4. Jupyter notebook - Effect of the CO2 uptake rate on C3 metabolism](notebooks/2019-05-06-mb-genC3-CO2-Effect.ipynb)   [![Binder](https://mybinder.org/badge_logo.svg)](https://mybinder.org/v2/gh/ma-blaetke/CBM_C3_C4_Metabolism/master?filepath=notebooks%2F2019-05-06-mb-genC3-CO2-Effect.ipynb)
- [Figure 1–source data 5. Jupyter notebook - Effect of the PPFD on C3 metabolism](notebooks/2019-05-06-mb-genC3-Light-Effect.ipynb).  [![Binder](https://mybinder.org/badge_logo.svg)](https://mybinder.org/v2/gh/ma-blaetke/CBM_C3_C4_Metabolism/master?filepath=notebooks%2F2019-05-06-mb-genC3-Light-Effect.ipynb)
- [Figure 1–source data 6. Jupyter notebook - Input, output fluxes and flux of energy in C3 metabolism](notebooks/2019-05-06-mb-genC3.ipynb)   [![Binder](https://mybinder.org/badge_logo.svg)](https://mybinder.org/v2/gh/ma-blaetke/CBM_C3_C4_Metabolism/master?filepath=notebooks%2F2019-05-06-mb-genC3.ipynb)

### Figure 3
- [Figure 3–source data 1. Jupyter notebook - Analysing the effect of oxygenation : carboxylation ratio on the emergence of the C4 cycle](notebooks/2019-05-06-mb-genC4-Decarb-Oxy-Ratio-Effect.ipynb)   [![Binder](https://mybinder.org/badge_logo.svg)](https://mybinder.org/v2/gh/ma-blaetke/CBM_C3_C4_Metabolism/master?filepath=notebooks%2F2019-05-06-mb-genC4-Decarb-Oxy-Ratio-Effect.ipynb)

### Figure 4
- [Figure 4–source data 1. Jupyter notebook - Effect of C4 mode on the emergence of the C4 cycle](notebooks/2019-05-06-mb-genC4-C4-mode.ipynb)   [![Binder](https://mybinder.org/badge_logo.svg)](https://mybinder.org/v2/gh/ma-blaetke/CBM_C3_C4_Metabolism/master?filepath=notebooks%2F2019-05-06-mb-genC4-C4-mode.ipynb)


### Figure 6
- [Figure 6–source data 1. Jupyter notebook - Effect of light on the C4 mode](notebooks/2019-05-06-mb-genC4-Light-Effect.ipynb)   [![Binder](https://mybinder.org/badge_logo.svg)](https://mybinder.org/v2/gh/ma-blaetke/CBM_C3_C4_Metabolism/master?filepath=notebooks%2F2019-05-06-mb-genC4-Light-Effect.ipynb)
- [Figure 6–source data 2. Jupyter notebook - Effect of NO3 – limitation on the C4 mode](notebooks/2019-05-06-mb-genC4-N-Limitation-Effect.ipynb)   [![Binder](https://mybinder.org/badge_logo.svg)](https://mybinder.org/v2/gh/ma-blaetke/CBM_C3_C4_Metabolism/master?filepath=notebooks%2F2019-05-06-mb-genC4-N-Limitation-Effect.ipynb)
- [Figure 6–source data 3. Jupyter notebook - Effect of H2O limitation on the C4 mode](notebooks/2019-05-06-mb-genC4-H2O-Limitation.ipynb)   [![Binder](https://mybinder.org/badge_logo.svg)](https://mybinder.org/v2/gh/ma-blaetke/CBM_C3_C4_Metabolism/master?filepath=notebooks%2F2019-05-06-mb-genC4-H2O-Limitation.ipynb)
- [Figure 6–source data 4. Jupyter notebook - Effect of CO2 limitation on the C4 mode](notebooks/2019-05-06-mb-genC4-CO2-Limitation.ipynb)   [![Binder](https://mybinder.org/badge_logo.svg)](https://mybinder.org/v2/gh/ma-blaetke/CBM_C3_C4_Metabolism/master?filepath=notebooks%2F2019-05-06-mb-genC4-CO2-Limitation.ipynb)
- [Figure 6–source data 5. Jupyter notebook - Effect of malate : asparte transport ratio on the C4 mode](notebooks/2019-05-06-mb-genC4-Mal_Asp-Ratio-Effect.ipynb)   [![Binder](https://mybinder.org/badge_logo.svg)](https://mybinder.org/v2/gh/ma-blaetke/CBM_C3_C4_Metabolism/master?filepath=notebooks%2F2019-05-06-mb-genC4-Mal_Asp-Ratio-Effect.ipynb)
