## Supplementary material for "Evolution of C4 photosynthesis predicted by constraint-based modelling": __init__.py

```
from __future__ import print_function
    
#cobra
global cobra
import cobra

#pandas
global pd
import pandas as pd


#numpy
global np
import numpy as np

#plotly
global ply, offline, download_plotlyjs, init_notebook_mode, iplot
import plotly as ply 
import plotly.offline as offline
from plotly.offline import download_plotlyjs, init_notebook_mode, iplot

#GO
global go
import plotly.graph_objs as go

#time
global sleep
from time import sleep

#os
global os
import os

#colorlover
global cl
import colorlover as cl

#operator
global operator
import operator

#escher
global Builder
from escher import Builder

#IPython
global display, HTML
from IPython.display import display
from IPython.core.display import HTML

#re
global re
import re

#myfunctions & constants
global inf
inf = float(1e6)

global init_notebook
from init_notebook import init_notebook

global load_sbml_model
from load_sbml_model import load_sbml_model

global add_rxn, set_bounds, set_fixed_flux, set_fixed_flux_ratio
from manipulate_model import add_rxn, set_bounds, set_fixed_flux, set_fixed_flux_ratio

global create_bar_plot_met, create_bar_plot_rxn, get_fluxes_by_metabolite, plot_transport, plot_transport_fva, create_scatter_plot_rxn_c3, create_scatter_plot_met_c3, create_bar_plot_met_c3
from view_results import create_bar_plot_met, create_bar_plot_rxn, get_fluxes_by_metabolite, plot_transport, plot_transport_fva, create_scatter_plot_rxn_c3, create_scatter_plot_met_c3, create_bar_plot_met_c3

global save_fba_to_excel, save_fva_to_excel, save_to_json
from save_results import save_fba_to_excel, save_fva_to_excel, save_to_json
```
