## Supplementary material for "Evolution of C4 photosynthesis predicted by constraint-based modelling": init_notebook.py

```
from init_fba import os, pd, init_notebook_mode

def init_notebook(theNotebook):
    
    #pandas
    pd.set_option('display.max_columns', 1000)
    pd.set_option('display.width', 10000)
    pd.options.display.max_colwidth = 500
    pd.set_option('display.max_rows', 1000)
    
    #init notebook mode for plotly
    init_notebook_mode(connected=True)
    
    #create folder to store results
    directory = theNotebook
    cwd = os.getcwd()
    if not os.path.exists(theNotebook):
        os.makedirs(theNotebook)
    if not os.path.exists(theNotebook+'/excel'):
        os.makedirs(theNotebook+'/excel')
    if not os.path.exists(theNotebook+'/figures'):
        os.makedirs(theNotebook+'/figures')
    if not os.path.exists(theNotebook+'/json'):
        os.makedirs(theNotebook+'/json')
    
    
```
