## Supplementary material for "Evolution of C4 photosynthesis predicted by constraint-based modelling": manipulate_model.py

```
def add_rxn(name, D_mets, model, rev=True):
    r_name = name
    r_obj = cobra.Reaction(rname)
    r_obj.name = r_name
    r_obj.id = r_name
    model.add_reaction(r_obj)
    r_obj.add_metabolites(D_mets)
    r_obj.objective_coefficient = 0
    r_obj.bounds = (-inf,inf) if rev else (0,inf)

def set_fixed_flux(r_id, val, model):
    r_obj = model.reactions.get_by_id(r_id)
    r_obj.bounds = (val,val)
    
def set_bounds(r_id, val_tuple, model):
    r_obj = model.reactions.get_by_id(r_id)
    r_obj.bounds = val_tuple
    
def set_fixed_flux_ratio(r_dict, model):
    if len(r_dict) == 2:
        r_id1 = r_dict.keys()[0]
        r_obj1 = model.reactions.get_by_id(r_id1)
        r_v1 = r_dict.values()[0]
        r_id2 = r_dict.keys()[1]
        r_obj2 = model.reactions.get_by_id(r_id2)
        r_v2 = r_dict.values()[1]
        const = model.problem.Constraint(r_v1 * r_obj2.flux_expression - r_v2 * r_obj1.flux_expression, lb = 0, ub = 0)
        model.add_cons_vars(const)
        return const
```
