## Supplementary material for "Evolution of C4 photosynthesis predicted by constraint-based modelling": save_results.py

```
from init_fba import pd, inf, re

def save_to_json(name, flux,theNotebook):
        js = flux.to_json()
        fp = open(theNotebook+'/json/'+name+'.json', 'w')
        fp.write(js)
        fp.close()

def save_fba_to_excel(c3_model, c4_model, D_fba, D_exp, theNotebook, file_name='pfba_results'):

    writer = pd.ExcelWriter(theNotebook+'/excel/'+file_name+'.xlsx')
    
    L_r_transport = []
    for r_obj in c4_model.reactions:
        if r_obj.id[0:3] == '[MB]':
            L_r_transport.append(r_obj.id)
        
    
    for exp in D_exp:
        DF_fba = pd.DataFrame(index=[r_obj.id for r_obj in c3_model.reactions]+L_r_transport,columns=['rxn','lb','ub','MC-Flux','BC-Flux',])
        for r_obj in c3_model.reactions:
            r_id = r_obj.id
            r_M_flux = D_fba[exp]['[M]_'+r_id]
            r_B_flux = D_fba[exp]['[B]_'+r_id]
            r_lb = '0' if r_obj.lower_bound == 0. else '-inf' if r_obj.lower_bound == -inf else str(r_obj.lower_bound)
            r_ub = '0' if r_obj.upper_bound == 0. else 'inf' if r_obj.upper_bound == inf else str(r_obj.lower_bound) 
            DF_fba.loc[r_id] = [r_obj.reaction,r_lb,r_ub,r_M_flux,r_B_flux]
        for r_id in L_r_transport:
            r_obj = c4_model.reactions.get_by_id(r_id)
            r_MB_flux = D_fba[exp][r_id]
            r_lb = '0' if r_obj.lower_bound == 0. else '-inf' if r_obj.lower_bound == -inf else str(r_obj.lower_bound)
            r_ub = '0' if r_obj.upper_bound == 0. else 'inf' if r_obj.upper_bound == inf else str(r_obj.lower_bound) 
            DF_fba.loc[r_id] = [r_obj.reaction,r_lb,r_ub,r_MB_flux, float('nan')]
        sheet = re.sub(r"[\[,:,*,?, \],\\,/]",'_', D_exp[exp])
        DF_fba.to_excel(writer,sheet)
    writer.save()
    
def save_fva_to_excel(c3_model, c4_model, D_fva, D_exp, theNotebook, file_name='fva_results'):

    writer = pd.ExcelWriter(theNotebook+'/excel/'+file_name+'.xlsx')
    
    L_r_transport = []
    for r_obj in c4_model.reactions:
        if r_obj.id[0:3] == '[MB]':
            L_r_transport.append(r_obj.id)
    
    for exp in D_exp:
        DF_fva = pd.DataFrame(index=[r_obj.id for r_obj in c3_model.reactions]+L_r_transport,columns=['rxn','lb','ub','MC-min','MC-max','BC-min','BC-max',])
        for r_obj in c3_model.reactions:
            r_id = r_obj.id
            r_M_max = D_fva[exp].get_value('[M]_'+r_id,'maximum')
            r_M_min = D_fva[exp].get_value('[M]_'+r_id,'minimum')
            r_B_max = D_fva[exp].get_value('[B]_'+r_id,'maximum')
            r_B_min = D_fva[exp].get_value('[B]_'+r_id,'minimum')
            r_lb = '0' if r_obj.lower_bound == 0. else '-inf' if r_obj.lower_bound == -inf else str(r_obj.lower_bound)
            r_ub = '0' if r_obj.upper_bound == 0. else 'inf' if r_obj.upper_bound == inf else str(r_obj.lower_bound)        
            DF_fva.loc[r_id] = [r_obj.reaction,r_lb,r_ub,r_M_min,r_M_max,r_B_min,r_B_max]
        for r_id in L_r_transport:
            r_obj = c4_model.reactions.get_by_id(r_id)
            r_MB_max = D_fva[exp].get_value(r_id,'maximum')
            r_MB_min = D_fva[exp].get_value(r_id,'minimum')
            r_lb = '0' if r_obj.lower_bound == 0. else '-inf' if r_obj.lower_bound == -inf else str(r_obj.lower_bound)
            r_ub = '0' if r_obj.upper_bound == 0. else 'inf' if r_obj.upper_bound == inf else str(r_obj.lower_bound)
            DF_fva.loc[r_id] = [r_obj.reaction,r_lb,r_ub,r_MB_min,r_MB_max, float('nan'), float('nan')]
        sheet = re.sub(r"[\[,:,*,?, \],\\,/]", '_', D_exp[exp])
        DF_fva.to_excel(writer,sheet)
    writer.save()
```
