## Supplementary material for "Evolution of C4 photosynthesis predicted by constraint-based modelling": view_results.py

```
# -*- coding: utf-8 -*-

from init_fba import go, iplot, sleep, cl, pd, np, operator, ply

def create_scatter_plot_rxn_c3(D_fba, L_r, title, xaxis_title, save_fig = False):
    data = []

    for r_id in L_r:
        trace = go.Scatter(y = [D_fba[exp][r_id] for exp in sorted(D_fba.keys())],
                           x = sorted(D_fba.keys()),
                           name = r_id,
                           mode = 'lines+markers',
                          )

        data.append(trace)

    layout = go.Layout(title = title,
                       yaxis = dict(title = 'Flux [µmol/s/m2]'),
                       xaxis = dict(title = xaxis_title),
                       width = 500,
                      )

    fig = go.Figure(data=data, layout=layout)
    
    if save_fig:
        iplot(fig,filename=title,image='svg',image_width=500,image_height=500)
        sleep(5)
    else:
        iplot(fig)
        
def create_scatter_plot_met_c3(model, D_fba, L_r, m_id, title, xaxis_title, save_fig = False):
    data = []

    for r_id in L_r:
        r_obj = model.reactions.get_by_id(r_id)
        m_v = r_obj.get_coefficient(m_id)
        trace = go.Scatter(y = [abs(m_v)*D_fba[exp][r_id] for exp in sorted(D_fba.keys())],
                           x = sorted(D_fba.keys()),
                           name = r_id,
                           mode = 'lines+markers',
                          )

        data.append(trace)

    layout = go.Layout(title = title,
                       yaxis = dict(title = 'Flux [µmol/s/m2]'),
                       xaxis = dict(title = xaxis_title),
                       width = 500,
                      )

    fig = go.Figure(data=data, layout=layout)
    
    if save_fig:
        iplot(fig,filename=title,image='svg',image_width=500,image_height=500)
        sleep(5)
    else:
        iplot(fig)

def create_bar_plot_met_c3(model, D_fba, L_r, m_id, title, xaxis_title, save_fig = False):
    data = []

    for r_id in L_r:
        r_obj = model.reactions.get_by_id(r_id)
        m_v = r_obj.get_coefficient(m_id)
        trace = go.Bar(y = [abs(m_v)*D_fba[exp][r_id] for exp in sorted(D_fba.keys())],
                           x = sorted(D_fba.keys()),
                           name = r_id,
                          )

        data.append(trace)

    layout = go.Layout(title = title,
                       yaxis = dict(title = 'Flux [µmol/s/m2]'),
                       xaxis = dict(title = xaxis_title),
                       width = 500,
                        barmode='stack',
                      )

    fig = go.Figure(data=data, layout=layout)
    
    if save_fig:
        iplot(fig,filename=title,image='svg',image_width=500,image_height=500)
        sleep(5)
    else:
        iplot(fig)

def create_bar_plot_rxn(D_fba, D_exp, D_rxn, title, xaxis_title, stacked = False, y_max = None, c = False, save_fig = False, absolute = True):
    
    #Coloring of bars
    L_colM = cl.scales['5']['seq']['Oranges']
    L_colM = L_colM[::-1]
    L_colB = cl.scales['5']['seq']['Blues']
    L_colB = L_colB[::-1]
    
    
    if len(D_rxn) >= 5 or c:
        L_colM = cl.scales['11']['qual']['Paired']
        L_colB = cl.scales['11']['qual']['Paired']
        
    if isinstance(D_rxn, list):
        D_rxn = {r_id: r_id for r_id in D_rxn}
    
    D_rid_colM = {r_id2: L_colM[i] for i, r_id2 in enumerate([r_id1 for r_id1 in D_rxn if r_id1[1] == 'M'])}
    D_rid_colB = {r_id2: L_colB[i] for i, r_id2 in enumerate([r_id1 for r_id1 in D_rxn if r_id1[1] == 'B'])}
    D_rid_col = D_rid_colM.copy()
    D_rid_col.update(D_rid_colB)
    
    #Create bar plot
    data = []
    for r_id, r_name in sorted(D_rxn.items()):

        trace = go.Bar(name = r_name,
                       y = [abs(D_fba[x][r_id]) for x in sorted(D_fba.keys())] if absolute else [D_fba[x][r_id] for x in sorted(D_fba.keys())],
                       x = [D_exp[exp]  for exp in sorted(D_fba.keys())],
                       marker = {'color': D_rid_col[r_id] if D_rid_col else '#1f77b4'} 
                      )
        data.append(trace)

    layout = go.Layout(height = 400,
                       width = 600,
                       margin = dict(b  = 100),
                       barmode='stack' if stacked else 'group',
                       xaxis= {'title': xaxis_title, 'tickangle':45 if len(D_exp) >= 8 else 0},
                       title = title,
                       yaxis = { 'title': 'Flux [µmol/s/m2]','range': [0, y_max] if y_max else None,'autorange': False if  y_max else True}
                       )

    fig = go.Figure(data=data, layout=layout)
    
    if save_fig:
        iplot(fig,image='svg', filename=title,image_width=600, image_height=400)
        sleep(5)
    else:
        iplot(fig)

def create_bar_plot_met(D_fba, D_exp, D_rxn, m_id, title, xaxis_title, c3_model, stacked = False, y_max = None, c = False, save_fig = False):
    
    #Coloring of bars
    L_colM = cl.scales['5']['seq']['Oranges']
    L_colM = L_colM[::-1]
    L_colB = cl.scales['5']['seq']['Blues']
    L_colB = L_colB[::-1]
    
    
    if len(D_rxn) >= 5 or c:
        L_colM = cl.scales['11']['qual']['Paired']
        L_colB = cl.scales['11']['qual']['Paired']
        
    if isinstance(D_rxn, list):
        D_rxn = {r_id: r_id for r_id in D_rxn}
    
    D_rid_colM = {r_id2: L_colM[i] for i, r_id2 in enumerate([r_id1 for r_id1 in D_rxn if r_id1[1] == 'M'])}
    D_rid_colB = {r_id2: L_colB[i] for i, r_id2 in enumerate([r_id1 for r_id1 in D_rxn if r_id1[1] == 'B'])}
    D_rid_col = D_rid_colM.copy()
    D_rid_col.update(D_rid_colB)
    
    #Create bar plot
    data = []
    for r_id, r_name in sorted(D_rxn.items()):
        r_obj = c3_model.reactions.get_by_id(r_id[4:])
        m_v = r_obj.get_coefficient(m_id)
        trace = go.Bar(name = r_name,
                       y = [abs(m_v*D_fba[x][r_id]) for x in sorted(D_fba.keys())],
                       x = [D_exp[exp]  for exp in sorted(D_fba.keys())],
                       marker = {'color': D_rid_col[r_id] if D_rid_col else '#1f77b4'} 
                      )
        data.append(trace)

    layout = go.Layout(height = 400,
                       width = 600,
                       margin = dict(b  = 100),
                       barmode='stack' if stacked else 'group',
                       xaxis= {'title': xaxis_title, 'tickangle':45 if len(D_exp) >= 8 else 0},
                       title = title,
                       yaxis = {'title': 'Flux [µmol/s/m2]','range': [0, y_max] if y_max else None,'autorange': False if  y_max else True})

    fig = go.Figure(data=data, layout=layout)
    
    if save_fig:
        iplot(fig,image='svg', filename=title,image_width=500,image_height=500)
        sleep(5)
    else:
        iplot(fig)
        

def plot_transport_MB(D_fba, D_exp, L_r, xaxis_title, save_fig, cut_off):
    
    df = pd.DataFrame(index = [], columns= [D_exp[exp]  for exp in sorted(D_fba.keys())])
    
    yTickNames_MB = []
    
    D_rid_meanFlux_MB = {r_id:abs(np.mean([0]+[D_fba[x][r_id] for x in D_exp if D_fba[x][r_id] > 0])) for r_id in L_r if abs(np.mean([0]+[D_fba[x][r_id] for x in D_exp if D_fba[x][r_id] > 0])) > cut_off}
    D_rid_meanFlux_MB = sorted(D_rid_meanFlux_MB.items(), key=operator.itemgetter(1))
    
    for entry in D_rid_meanFlux_MB:
        r_id = entry[0]
        for x in D_exp:
            if D_fba[x][r_id] > 0:
                df.set_value(r_id, D_exp[x], abs(D_fba[x][r_id]) if not abs(D_fba[x][r_id]) < cut_off else float('nan') )
                if not r_id[5:-2] in yTickNames_MB:
                    yTickNames_MB.append(r_id[5:-2])
                        
    scl = cl.scales['9']['seq']['Reds']
    colorscale = [ [ float(i)/float(len(scl)-1), scl[i] ] for i in range(len(scl))]
    
    trace_MB = go.Heatmap(z=df.values.tolist(),
                          x=df.columns,
                          y=yTickNames_MB,
                          colorbar= {'title':'Flux [µmol/s/m2]', 'titleside':'right'},
                          colorscale = colorscale)
    
    layout_MB = go.Layout(width = 500,
                          margin = {'b':100},
                          yaxis = {'tickmode': 'array', 'tickvals': range(0,len(yTickNames_MB)), 'ticktext': yTickNames_MB, 'title': 'Transport Metabolites'},
                          xaxis= {'title':xaxis_title,'tickangle':45 if len(D_exp) >= 8 else 0},
                          title = 'Mesophyll ==> Bundlesheat Transport')
    
    data_MB=[trace_MB]
    
    fig_MB = go.Figure(data=data_MB, layout=layout_MB)
    if save_fig:
        iplot(fig_MB,image='svg', filename='transport_MB',image_width=500,image_height=500)
        sleep(5)
    else:
        iplot(fig_MB)
        
    
def plot_transport_BM(D_fba, D_exp, L_r, xaxis_title, save_fig, cut_off):
    
    df = pd.DataFrame(index = [], columns= [D_exp[exp]  for exp in sorted(D_fba.keys())])
    
    yTickNames_BM = []
        
    D_rid_meanFlux_BM = {r_id:abs(np.mean([0]+[D_fba[x][r_id] for x in D_exp if D_fba[x][r_id] < 0])) for r_id in L_r if abs(np.mean([0]+[D_fba[x][r_id] for x in D_exp if D_fba[x][r_id] < 0])) > cut_off}
    D_rid_meanFlux_BM = sorted(D_rid_meanFlux_BM.items(), key=operator.itemgetter(1))
    
    for entry in D_rid_meanFlux_BM:
        r_id = entry[0]
        for x in D_exp:
            if D_fba[x][r_id] < 0:
                df.set_value(r_id, D_exp[x], abs(D_fba[x][r_id]) if not abs(D_fba[x][r_id]) < cut_off else float('nan') )
                if not r_id[5:-2] in yTickNames_BM:
                    yTickNames_BM.append(r_id[5:-2])
    
    scl = cl.scales['9']['seq']['Blues']
    colorscale = [ [ float(i)/float(len(scl)-1), scl[i] ] for i in range(len(scl)) ]
    
    trace_BM = go.Heatmap(z=df.values.tolist(),
                          x=df.columns,
                          y=yTickNames_BM,
                          colorbar= {'title':'Flux [µmol/s/m2]', 'titleside':'right'},
                          colorscale = colorscale)
    
    layout_BM = go.Layout(width = 500,
                          margin = {'b':100},
                          yaxis = {'tickmode': 'array', 'tickvals': range(0,len(yTickNames_BM)), 'ticktext': yTickNames_BM, 'title': 'Transport Metabolites'},
                          xaxis= {'title':xaxis_title,'tickangle':45 if len(D_exp) >= 8 else 0},
                          title = 'Bundlesheat ==> Mesophyll Transport')
    
    data_BM=[trace_BM]
    
    fig_BM = go.Figure(data=data_BM, layout=layout_BM,)
    
    if save_fig:
        iplot(fig_BM,image='svg', filename='transport_BM',image_width=500,image_height=500)
        sleep(5)
    else:
        iplot(fig_BM)  
    

def plot_transport(D_fba, D_exp, L_r, xaxis_title, save_fig = False):
    
    cut_off = 0.1
    
    plot_transport_MB(D_fba, D_exp, L_r, xaxis_title, save_fig, cut_off)
    plot_transport_BM(D_fba, D_exp, L_r, xaxis_title, save_fig, cut_off)
        
def plot_transport_fva(c4_model, D_exp, D_pfva, D_pfba, L_r_org, flux_max, title, save_fig=False, L_r_index=None):
    
    L_colors = ply.colors.DEFAULT_PLOTLY_COLORS
    
    if len(D_exp) > len(L_colors):
        print('Too many experiments.')
    else:
        p_value = 0.1 #p value
        
        #prepare data for plotting
        D_r_flux_range = {}
        D_exp_r_FVA ={}
        data = []
        
        for exp in D_exp:
            D_exp_r_FVA[exp] = {}
            for r_id in L_r_org:
                D_FVA = {}
                r_obj = c4_model.reactions.get_by_id(r_id)
                D_FVA['max'] = D_pfva[exp].get_value(r_id,'maximum')
                D_FVA['min'] = D_pfva[exp].get_value(r_id,'minimum')
                D_FVA['flux'] = D_pfba[exp][r_id]
                
                #filter flux ranges of particular size (p_value % of maximum C transport rate)
                if abs(D_FVA['max']) > p_value * flux_max or abs(D_FVA['min']) > p_value * flux_max:
                    #determine maximum flux range over all exp for each reaction
                    if not r_id in D_r_flux_range: 
                        D_r_flux_range[r_id] = D_FVA['max'] - D_FVA['min']
                    else:
                        if D_r_flux_range[r_id] < (D_FVA['max'] - D_FVA['min']):
                            D_r_flux_range[r_id] = D_FVA['max'] - D_FVA['min']
                            
                D_exp_r_FVA[exp][r_id] = D_FVA
        
        #sort reactions by size of flux range
        L_r = [r_flux_range[0] for r_flux_range in sorted(D_r_flux_range.items(), key=operator.itemgetter(1), reverse= True)]
        
        create_plot = True
        
        if L_r_index:
            L_r_ex1 = list(set(L_r_index)-set(L_r))
            L_r_ex2 = list(set(L_r)-set(L_r_index))
            L_r_ex = L_r_ex1 + L_r_ex2
            if not L_r_ex:
                L_r = L_r_index
            else:
                print('Warning: L_r and L_r_index are not matching: %s' %L_r_ex)
                create_plot = False
                
        if create_plot:
                #set up x-values for each experiment
                L_x_axis = np.arange(1,len(L_r)*2+1,2)
                D_exp_x_axis = {exp: [] for exp in D_exp}
                for x in L_x_axis:
                    L_x_exp = np.linspace(x-0.3,x+0.3,len(D_exp))
                    for i_exp, exp in enumerate(D_exp.keys()):
                        D_exp_x_axis[exp].append(L_x_exp[i_exp]) 
                        
                #prepare trace for each experiment
                for i_exp, (exp,exp_name) in enumerate(D_exp.items()):
                    L_r_flux = [D_exp_r_FVA[exp][r_id]['flux'] for r_id in L_r]
                    L_r_min= [D_exp_r_FVA[exp][r_id]['flux'] - D_exp_r_FVA[exp][r_id]['min'] for r_id in L_r]
                    L_r_max = [D_exp_r_FVA[exp][r_id]['max'] - D_exp_r_FVA[exp][r_id]['flux'] for r_id in L_r]
                    
                    trace = go.Scatter(
                        x = D_exp_x_axis[exp],
                        y = L_r_flux,
                        error_y={'type':'data', 'symmetric':False, 'array':L_r_max, 'arrayminus':L_r_min},
                        marker = {'color' : L_colors[i_exp]},
                        name = exp_name,
                        mode = 'markers'
                    )
            
                    data.append(trace)
                
                #prepare layout
                layout = go.Layout(
                    xaxis = {'tickvals':L_x_axis, 'ticktext':[r_id[5:-2] for r_id in L_r], 'title':'Exchange Metabolites'},
                    #margin = {'b': 300},
                    title = title,
                    yaxis = {'title':'Flux [µmol/s/m2]'},
                    legend = {'orientation':'h', 'x':0, 'y':-0.15},
                    shapes = [
                            {
                                'type': 'rect',
                                'x0': -0.1,
                                'y0': -0.5,
                                'x1': -1.5,
                                'y1': -50,
                                'line': {
                                    'color': 'rgba(255, 255, 255, 1)',
                                    'width': 1,
                                },
                                'fillcolor': 'rgba(255, 255, 255, 1)',
                            },
                            {
                                'type': 'rect',
                                'x0': -0.1,
                                'y0': 0.5,
                                'x1': -1.5,
                                'y1': 50,
                                'line': {
                                    'color': 'rgba(255, 255, 255, 1)',
                                    'width': 1,
                                },
                                'fillcolor': 'rgba(255, 255, 255, 1)',
                            },
                    ],
                    annotations=[
                        dict(
                            x=-0.5,
                            y=25,
                            xref='x',
                            yref='y',
                            text='Mesophyll -> Bundle Sheath',
                            textangle= -90,
                            showarrow=False,
                            font=dict(
                                size=10,
                                color='#000000'
                            ),
                            align='center',
                        ),
                        dict(
                        x=-0.5,
                        y=-25,
                        xref='x',
                        yref='y',
                        text='Bundle Sheath -> Mesophyll',
                        textangle= -90,
                        showarrow=False,
                        font=dict(
                            size=10,
                            color='#000000'
                        ),
                        align='center',
        
                    )
                ]
                )
                
                #create figure
                fig = go.Figure(data=data, layout=layout)
                if save_fig:
                    iplot(fig,image='svg', filename='FVA')
                    sleep(5)
                else:
                    iplot(fig) 
    

def get_fluxes_by_metabolite(D_exp, D_fba, model, m_id, cell):
    if cell in ['M','B']:
        df = pd.DataFrame(columns=['rxn']+[D_exp[exp] for exp in D_exp])
        m_obj = model.metabolites.get_by_id('['+cell+']_'+m_id)
        for r_obj in m_obj.reactions:
            if not r_obj.id in df.index:
                df.loc[r_obj.id] = [r_obj.reaction] + [round(D_fba[exp][r_obj.id],3) for exp in D_exp]
        return df
    else:
        return 'Choose either M or B as cell type'
```
